# Conversion of a defensive toxin-antitoxin system into an offensive T6SS effector in Burkholderia

**DOI:** 10.1101/835710

**Authors:** Sunil Kumar Yadav, Ankita Magotra, Aiswarya Krishnan, Srayan Ghosh, Rahul Kumar, Joyati Das, Gopaljee Jha

## Abstract

Bacteria use various kinds of toxins to either inhibit the growth of co-habiting bacteria or when needed control their own growth. Here we report that Burkholderia and certain other bacteria have altered the potential defensive function of Tox-REase-5 domain containing toxins into offensive function. The *Burkholderia gladioli* strain NGJ1 encodes such toxins as type VI secretion system (T6SS) effectors (Tse) and potentially deploys them to kill co-habiting rice endophytic bacteria. Notably, the immunity (Tsi) proteins associated with Tse effectors demonstrate functional similarity with the antitoxin of type II toxin-antitoxin (TA) system. Genome analysis of diverse bacteria revealed that various Tse orthologs are either encoded as TA or T6SS effectors. In addition, potential evolutionary events associated with conversion of TA into T6SS effectors have been delineated. Our results indicate that the transposition of IS3 elements has led to the operonic fusion of certain T6SS related genes with TA genes resulting in their conversion into T6SS effectors. Such a genetic change has enabled bacteria to utilize novel toxins to precisely target co-habiting bacteria.

## Introduction

Under natural conditions, bacteria have to compete with co-habiting bacteria for available resources. This can lead to severe evolutionary pressure on bacteria to adopt strategies to limit the growth of other cohabiting microbes^1^. Several bacterial species use a specialized protein secretion system called the type VI secretion system (T6SS) to target co-habiting bacteria^2–6^. The T6SS is a syringe like apparatus composed of a base plate, a membrane complex (spanning the inner and outer membrane) and an inner tube, being wrapped in a sheath-like structure^7–10^. Hexamers of the HCP (hemolysin-coregulated protein) protein forms the inner tube of the T6SS apparatus^9,11^. A trimer of the VgrG (Valine-glycine repeat protein G) protein forms a spike like structure on the top of the inner tube^12^. Further, the PAAR (proline-alanine-alanine-arginine) repeat-containing protein binds to the distal end of the spike and forms a sharp pointed tip^13,14^. Contraction of the sheath enables the HCP-VgrG-PAAR protein complex to puncture the bacterial membrane and deliver various T6SS effectors into the extracellular environment or directly into the target bacterial cells^8,15,16^. The effectors can be encoded either as fused protein with the HCP/VgrG/PAAR proteins as an additional domain or they are non-covalently fused to HCP/VgrG/PAAR protein that are encoded as upstream ORF in the effector operon^12,15,17–20^. Association with HCP/VgrG/PAAR proteins/domains is essential for translocation of effectors. Recent studies have suggested that certain chaperone (DUF4123, DUF1795 and DUF2169) and co-chaperones are also required for T6SS mediated delivery of effectors^20–23^. Till date, diverse kinds of proteins including phospholipases (Tli), amidases (Tae), glucosaminidase (Tge), nucleases (Rhs proteins), DNases (Tde), peptidases, pore-forming toxins etc. have been identified as T6SS effectors from different bacteria^24–26^. These effectors demonstrate potent bactericidal activity by targeting the DNA/RNA/cell wall components of the prey bacterium^23–25^. However to protect self as well as sister cells from intoxication, bacteria have evolved cognate immunity proteins against each T6SS effector^25,27,28^. The effector-immunity pairs are encoded together in an operonic fashion and the immunity protein neutralizes toxic effect of the cognate effector via direct binding/interaction.

Beside T6SS effectors, bacteria encode various toxins as part of toxin-antitoxin (TA) system^29–31^. Similar to that of immunity proteins, the antitoxin of TA system bind to the cognate toxin and neutralize, to keep the cells protected. However, under certain environmental stress condition, the antitoxin gets degraded and thereby releases the toxin to exert antibacterial activity. This causes rapid growth arrest, formation of persistence and antibiotic resistant bacterial cells. However in certain kind of TA system (called type II TA system), besides directly neutralizing the toxin, the antitoxin alone or in complex with cognate toxin binds to the promoter of the TA operon and repress the expression of toxin-antitoxin genes^29,31^. The degradation of antitoxin, leads to release of toxin and also causes de-repression of toxin gene expression. Notably, the toxin and antitoxin ratio decides the fate of the cells.

In the present study, we demonstrate that a rice associated bacterium, *Burkholderia gladioli* strain NGJ1 utilizes two different T6SSs (here named as T6SS-1 and T6SS-2) to target a wide spectrum of co-habiting bacteria. Bioinformatics analysis indicates that the NGJ1 bacterium encodes fourteen different T6SS effectors and immunity proteins. Amongst them the restriction endonuclease (Tox-REase-5) domain containing effectors (here onward referred as Tse; type VI secreted effector) were quite noteworthy. The antitoxin (type VI secreted effector’s immunity; Tsi) associated with these effectors showed transcriptional repression activity. Genome organization and functional similarity suggest that the Tse effectors have potentially evolved from a bacterial TA system. The transposition of IS3 elements appears to have played a major role in this evolutionary adaptation which has enabled the bacteria to re-rationalize the function of these toxins as extracellular weapons to kill other bacteria.

## Results

### *B. gladioli* strain NGJ1 uses two different type VI secretion systems (T6SS) for antibacterial activity

We observed that *B. gladioli* strain NGJ1 demonstrates strong antibacterial activity against several rice endophytic bacteria as well as *Escherichia coli* and *Agrobacterium tumefaciens* (Supplementary Fig. 1 and Supplementary Table 1). Genome analysis revealed NGJ1 to contain two different T6SS apparatus encoding gene clusters, here named as T6SS-1 (Burkholderia genome database locus id: *ACI79_RS13910*-*ACI79_RS13980)* and T6SS-2 (Burkholderia genome database locus id: *ACI79_RS29585*-*ACI79_RS29710)* (Fig. 1a). Through plasmid integration, we disrupted one important gene from each of the clusters to obtain ΔT6SS-1 (*VipA* gene was disrupted) and ΔT6SS-2 mutants (*ImpE;* gene was disrupted). The western blot analysis revealed that wild type NGJ1 is able to secrete HCP-1 (associated with T6SS-1 cluster) and HCP-2 (Associated with T6SS-2 cluster) proteins into the extracellular milieu. This suggests that both of the T6SSs are functional. However, ΔT6SS-1 and ΔT6SS-2 mutants were defective in secreting their respective HCP proteins (Fig. 1b and c).

**Fig. 1.**
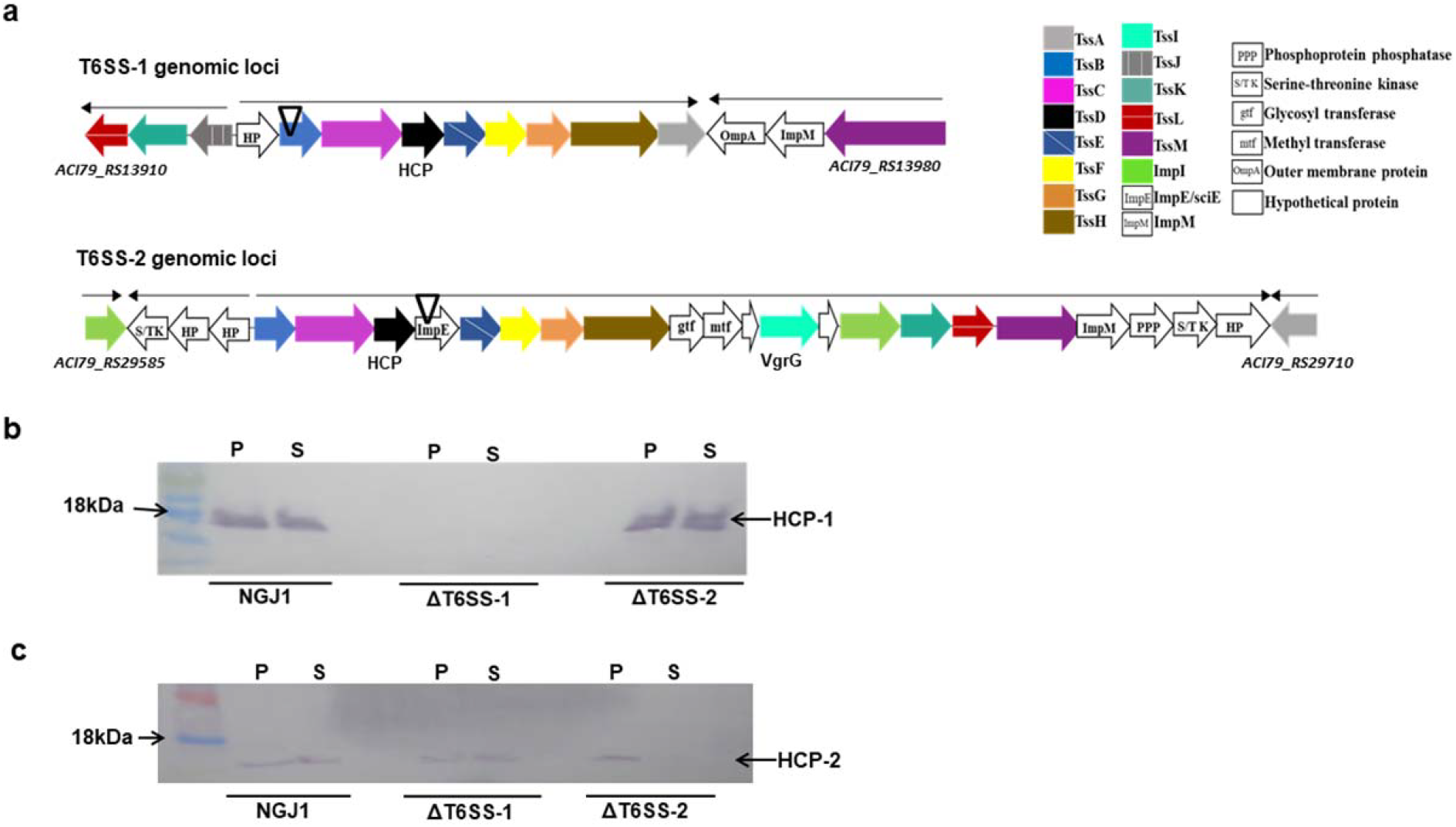
*B. gladioli* strain NGJ1 encodes two different T6SS apparatus. (a) Schematic representation of genomic organization of two different T6SS apparatus encoding operons (named as T6SS-1 and T6SS-2) in NGJ1 genome. Common components of the apparatus are color coded while unique components are indicated by their particular names in empty boxes. (b) Secretion profile of HCP protein in different NGJ1 strains. The total proteins from cell-free supernatant (S) as well as whole-cell lysates (P) of different strains were immunoblotted using HCP-1 (associated with T6SS-1) and HCP-2 (associated with T6SS-2) specific antibodies. Both the HCP proteins were detected in the supernatant of NGJ1 culture, while only HCP-2 was detected in ΔT6SS-1 and HCP-1 in ΔT6SS-2, culture supernatants. Similar results were obtained in at least two independent experiments.

We further tested the antibacterial ability of ΔT6SS-1 and ΔT6SS-2 mutants. The disruption of either of T6SS-1 or T6SS-2 had compromised the antibacterial activity of NGJ1 on most of the tested bacteria (Supplementary Fig. 1 and Supplementary Table 1). However in few cases, ΔT6SS-1 had lost antibacterial activity but ΔT6SS-2 mutant remained proficient in killing them. This suggests that NGJ1 target these bacteria in a T6SS-1 dependent manner.

### *B. gladioli* strain NGJ1 harbors diverse antibacterial T6SS effectors

Using computational analysis, we identified 14 different T6SS effector operons in the NGJ1 genome (Supplementary Fig. 2). Besides encoding effector proteins, each operon also encoded cognate immunity protein and certain carrier (VgrG and/or PAAR) as well as chaperone (DUF4123/ DUF1795) proteins which potentially assist in T6SS mediated delivery of effectors (Supplementary Fig. 2). The putative functions of various T6SS effectors of NGJ1 are summarized in Supplementary Table 2. Several effectors were chosen for ectopic expression and they were found to be lethal for the recombinant *E. coli* cells. In most of the cases, co-expression of cognate immunity proteins protected the cells from effector mediated killing (Supplementary Fig. 3). Interestingly, co-expression of Tse effectors (17tse/38tse) and their respective Tsi immunity (17tsi/38tsi) proteins using two different plasmids (pET23b:effector + pET28a:immunity) failed to protect the *E. coli* cells from effector mediated killing (Fig. 2a and b). However, when transcriptionally fused effector-immunity (17tsei/38tsei) proteins were expressed using a single plasmid, the recombinant *E. coli* cells were protected (Fig. 2c).

**Fig. 2.**
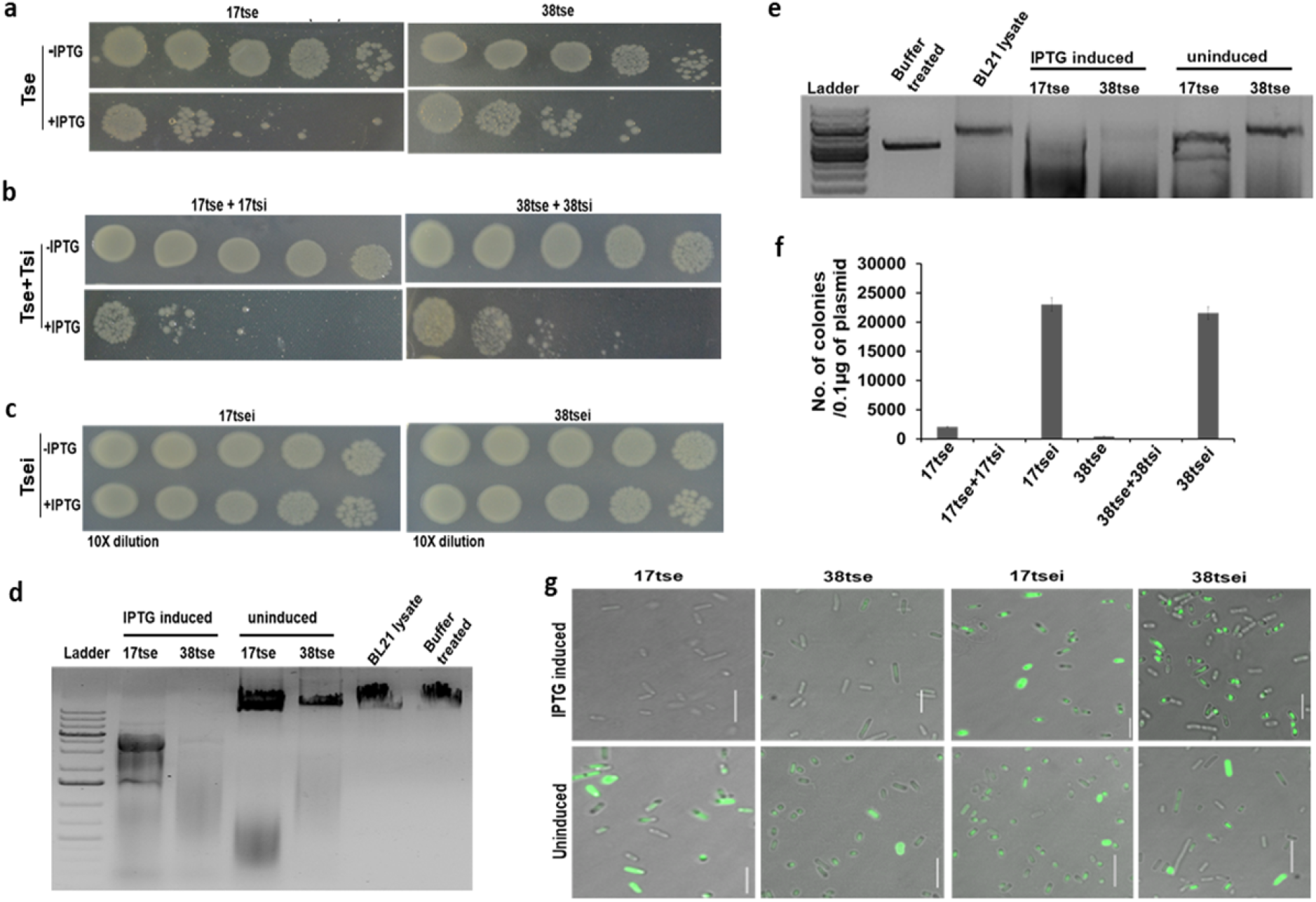
The Tse effectors of NGJ1 exhibit anti-bacterial activity. (a) IPTG induced expression of Tse protein inhibited the growth of recombinant *E. coli* (BL21) cells. (b) Co-expression of the immunity protein (Tsi) and the cognate Tse effector protein using two different plasmids (referred as Tse + Tsi) could not protect the cells from effector mediated killing. (c) Coupled-expression of transcriptionally fused Tse and Tsi proteins using a single plasmid (Tsei) protected the cells. (d) Treatment with crude preparation of Tse proteins caused the degradation of lambda DNA and (e) bacterial plasmid (*EcoR*I digested pET23b). (f) Graph showing number of *E. coli* (DH5α) cells obtained upon transformation of plasmids that were isolated from various Tse/ Tse+Tsi/ Tsei protein expressing *E. coli* (BL21) cells. Graph shows mean values ± s.d. (g) Fluorescence microscopic images of SYTOX-green (nucleic acid staining dye) stained *E. coli* (BL21) cells that express Tse/ Tsei proteins. Lack of staining in the effector expressing cells suggested DNA degradation while proper staining in the transcriptionally fused effector-immunity (Tsei) expressing cells suggested intact DNA (scale bar = 10µm). Similar results were obtained in at least three independent experiments.

**Fig. 3.**
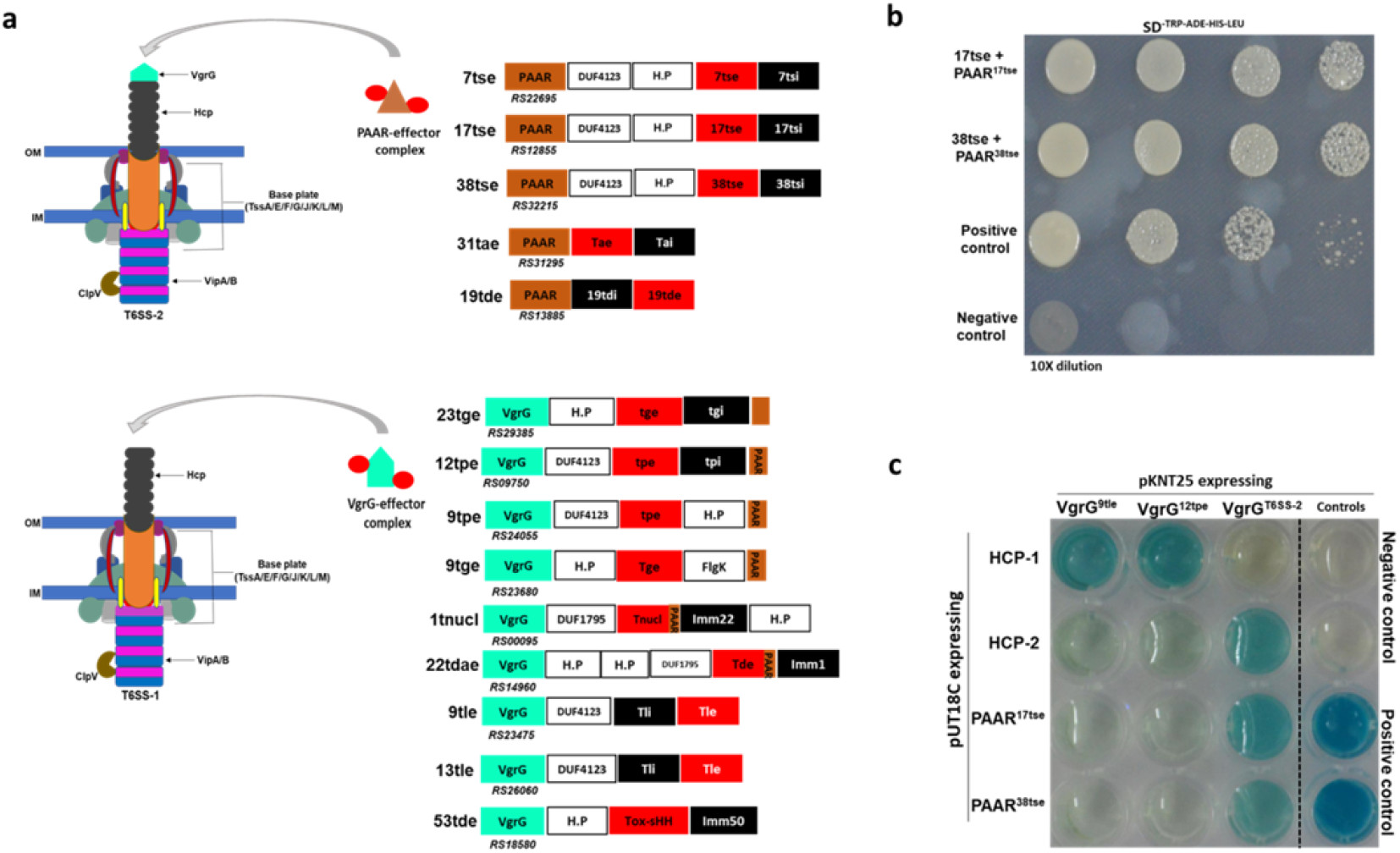
The Tse effectors are potentially secreted by T6SS-2 of NGJ1. (a) Schematic representation of potential mode of secretion of various T6SS effectors. The PAAR containing effectors are anticipated to be delivered by T6SS-2 while the VgrG containing effectors are thought to be delivered by T6SS-1. (b) Yeast two hybrid assay demonstrating positive interaction of Tse (17tse and 38tse) effector with the corresponding PAAR protein (PAAR^Tse^) encoded in their operon. Interaction between p53 and SV40 large T-antigen (T) proteins were used as positive control. The pGBKT7 and pGADT7 (empty vectors) were used as negative control. (c) Bacterial two hybrid assay demonstrating interaction between various components of T6SS apparatus and effector operons. Interaction between T25-Zip and T18-Zip was used as positive control while pKNT25 and pUT18C (empty vectors) were used as negative control. Appearance of blue color suggested positive interaction while absence of color suggested no interaction. Interaction of PAAR^Tse^ with the VgrG^T6SS-2^ and HCP-2 with VgrG^T6SS-2^ suggested the delivery of Tse effectors through T6SS-2. Similar results were obtained in at least three independent experiments.

### The Tse effectors of NGJ1 have endonuclease activity

NCBI Conserved Domain analysis revealed presence of a conserved restriction endonuclease-5 (Tox-REase-5) domain at the C-terminus of the Tse effectors (Supplementary Fig. 4a). This suggested that they might have endonuclease activity. Treatment with the crude preparation of Tse protein (17tse/38tse) caused degradation of lambda DNA (Fig. 2d) as well as bacterial plasmid (Fig. 2e). Moreover, the plasmid isolated from effector (17tse/38tse) or effector and cognate immunity (17tse + 17tsi/ 38tse + 38tsi) expressing cells was potentially fragmented as only limited number of bacterial colonies was obtained when these plasmids were transformed into *E. coli* (Fig. 2f). On the other hand, the plasmid isolated from transcriptionally fused effector-immunity (17tsei/38tsei) expressing cells yielded a large number of colonies upon transformation in *E. coli*.

**Fig. 4.**
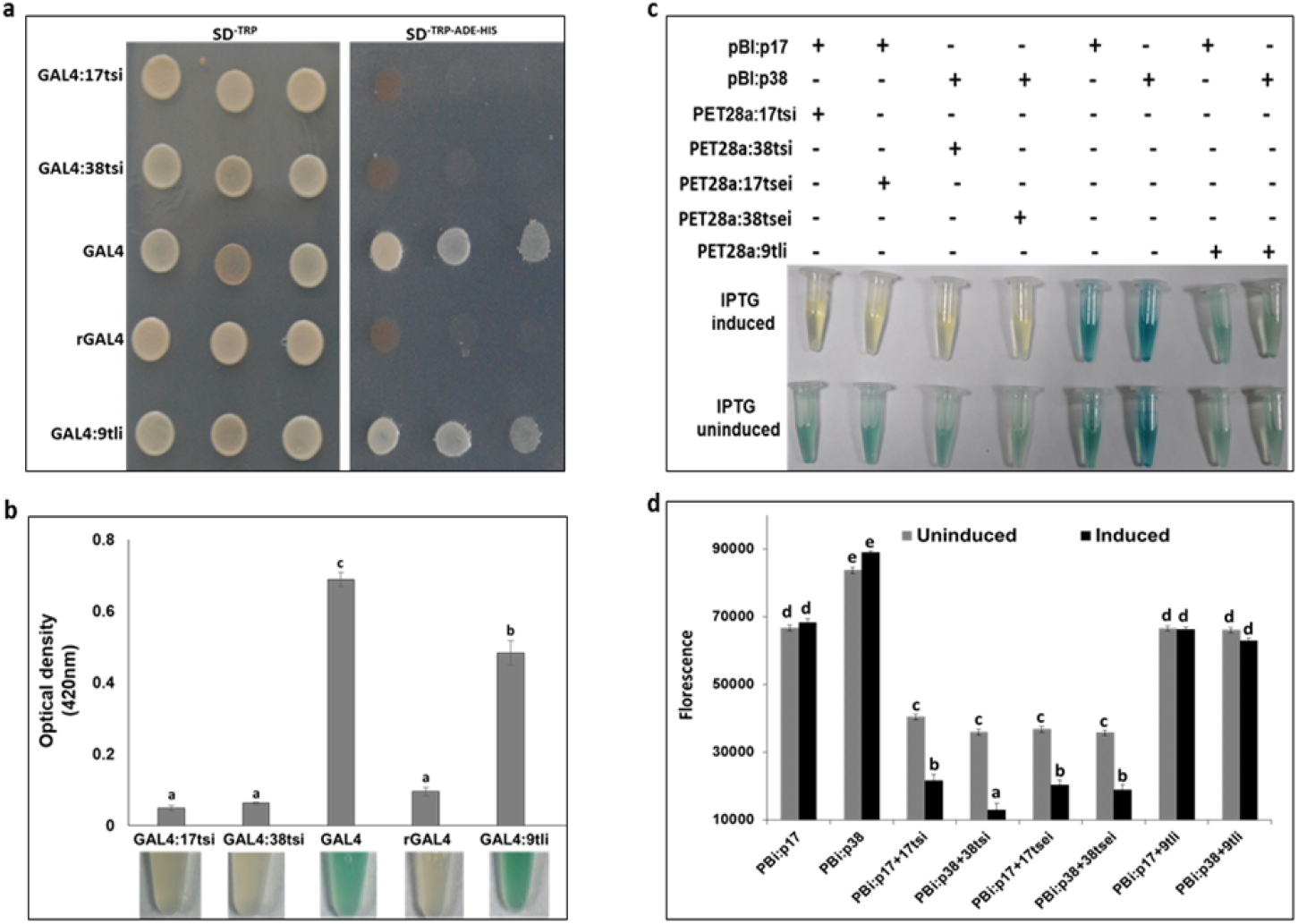
The Tsi immunity proteins of NGJ1 demonstrate transcriptional repressor activity. (a) Trans-repression assay in *S. cerevisiae* (yeast) wherein cells expressing GAL4 fused with Tsi immunity proteins (GAL4:17tsi/GAL4:38tsi) were unable to express reporter gene. The recombinant cells showed auxotrophy to ADE (adenine) and HIS (histidine). On the other hand, cells expressing native GAL4 or GAL4 fused with 9tli immunity protein (GAL4:9tli) were able to drive the expression of reporter genes. The rGAL4 (lacking activation domain) was used as negative control. (b) Expression of β-galactosidase reporter gene in recombinant yeast cells as revealed by X-gal (5-bromo-4-chloro-3-indolyl-β-D-galactopyranoside) coloration and calorimetric estimation. (c) The Tsi immunity proteins can repress their own promoter. The promoter of 17tse and 38tse encoding operon could drive the expression of reporter gene (*GUS;* β-glucuronidase) in *E. coli* as revealed by appearance of blue color. However, the *GUS* expression was repressed in presence of immunity (17tsi/ 38tsi) or effector-immunity (17tsei/ 38tsei) proteins. In presence of 9tli immunity protein, the *GUS* expression was observed. (d) Fluorometric quantification of GUS protein by MUG (4-methylumbelliferyl β-D-glucuronide) assay. Values with different letters are significantly different at P<0.001 (estimated using one-way ANOVA). Graphs show mean values ± standard deviation. Similar results were obtained in at least three independent experiments.

Next, we visualised the degradation of nucleic acid in recombinant *E. coli* cells using SYTOX Green dye. Lack of staining was observed in cells that express effector (17tse/38tse) or co-express effector and immunity proteins (17tse+17tsi/38tse+38tsi) on separate plasmids. However, intense staining was observed in cells that express transcriptionally fused effector-immunity pairs (17tsei/38tsei) (Fig. 2g and Supplementary Fig. 5). Taken together, these results indicate that Tse effectors can function as endonuclease.

**Fig. 5.**
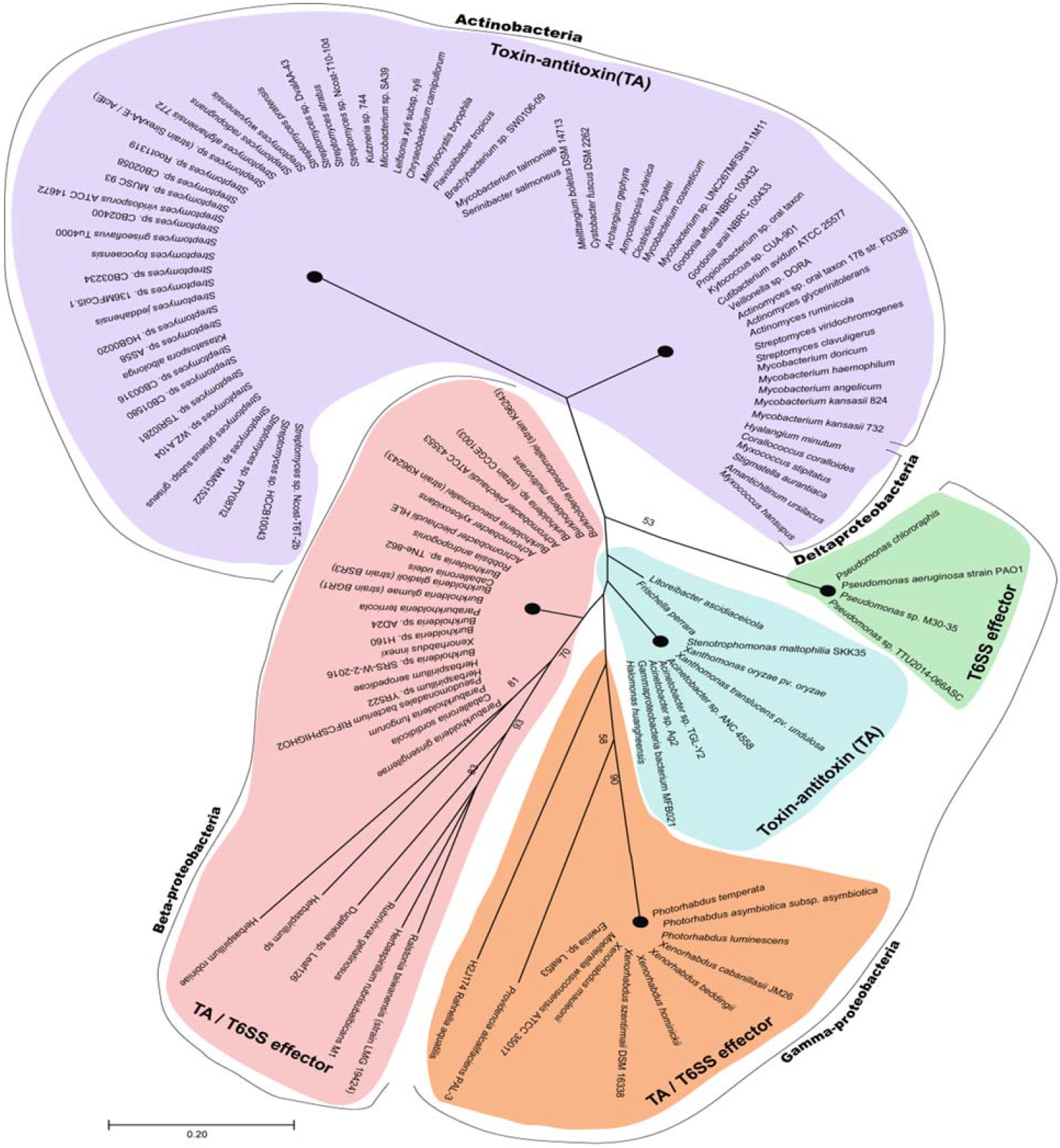
The Tse orthologs are conserved in different gram-negative bacteria. Pfam database search revealed orthologs of Tse proteins (n=199) to be present in different bacterial species. Phylogenetic tree of different Tse orthologs constructed using neighbor-joining algorithm is reflected. The bootstrap value is depicted at each node and the node with bootstrap value less than 50% has been condensed. The tree is drawn to scale where branch lengths denote the evolutionary distance. Based upon the presence of Tse orthologs as T6SS effector or TA or both, the clades are color coded. The bacterial classification is denoted in the outermost bar line.

### The Tse effectors are potentially secreted through T6SS-2 of NGJ1

The VgrG and PAAR proteins play important roles in secretion of T6SS effectors ^13,14,16,23^. In NGJ1, we observed either VgrG or PAAR are encoded in the upstream region of various effector operons, indicating that they might act as a carrier for their T6SS mediated delivery (Fig. 3a and Supplementary Fig. 2). The VgrG and PAAR are known to be encoded in some but not all T6SS apparatus encoding gene clusters^23,32,33^.As shown in Fig 1a, the T6SS-2 gene cluster encodes a VgrG protein but lacks PAAR protein, while the T6SS-1 cluster lacks both VgrG and PAAR. Considering this, we hypothesized that effectors which have VgrG (VgrG^effector^) encoded in the same operon might be secreted through T6SS-1 while those which are not co-encoded with VgrG would perforce have to be secreted through T6SS-2 (Fig. 3a).

The Tse effector encoding operons of NGJ1 lacks VgrG but encode PAAR (PAAR^Tse^) as upstream ORF. Therefore they would employ the PAAR as a carrier for their T6SS mediated delivery. This was confirmed by yeast two hybrid assays which demonstrated interaction of the PAAR^Tse^ with the cognate Tse proteins (Fig. 3b). Furthermore, the bacterial two hybrid assay revealed that the PAAR^Tse^ has strong binding affinity with the VgrG of T6SS-2 apparatus (VgrG^T6SS-2^) but not with the VgrG of effector operons (VgrG^9tle^/ VgrG^12tpe^). This suggested that the Tse effectors are secreted through T6SS-2 via interaction of PAAR^Tse^ with VgrG^T6SS-2^. In this regard, we observed that the VgrG^T6SS-2^ has specific binding affinity with HCP-2, but not with the HCP-1 (Fig. 3c). On the other hand, VgrG^9tle^ and VgrG^12tpe^ demonstrated specific binding affinity with HCP-1 but not with HCP-2, suggesting them to be secreted through T6SS-1 (Fig. 3c).

### Antitoxins associated with the Tse proteins of NGJ1 demonstrate transcriptional repressor activity

Bioinformatics analysis revealed that the immunity proteins (Tsi) associated with the Tse effectors contains an Imm52 domain (Supplementary Fig. 4b). Phylogenetic analysis suggested them to function as bacterial LysR family transcriptional regulators (Supplementary Fig. 6). They also shared structural similarity with a CodY family of pleotropic transcriptional repressor (Supplementary Table 3). Considering the above, we investigated the possible transcriptional repressor activity of 17tsi and 38tsi immunity proteins in a yeast trans-repression assay^34,35^. When GAL4 was transcriptionally fused with 17tsi/ 38tsi, it caused repression of reporter gene (*lacZ*, *HIS3* and *ADE2*) expression in yeast. The recombinant cells showed auxotrophy to histidine as well as adenine (Fig. 4a and b). On the other hand, GAL4 transcriptionally fused with immunity protein (9tli) of lipase effector (which does not have transcription repression activity) induced the expression of reporter genes (Fig. 4a and b).

**Fig. 6.**
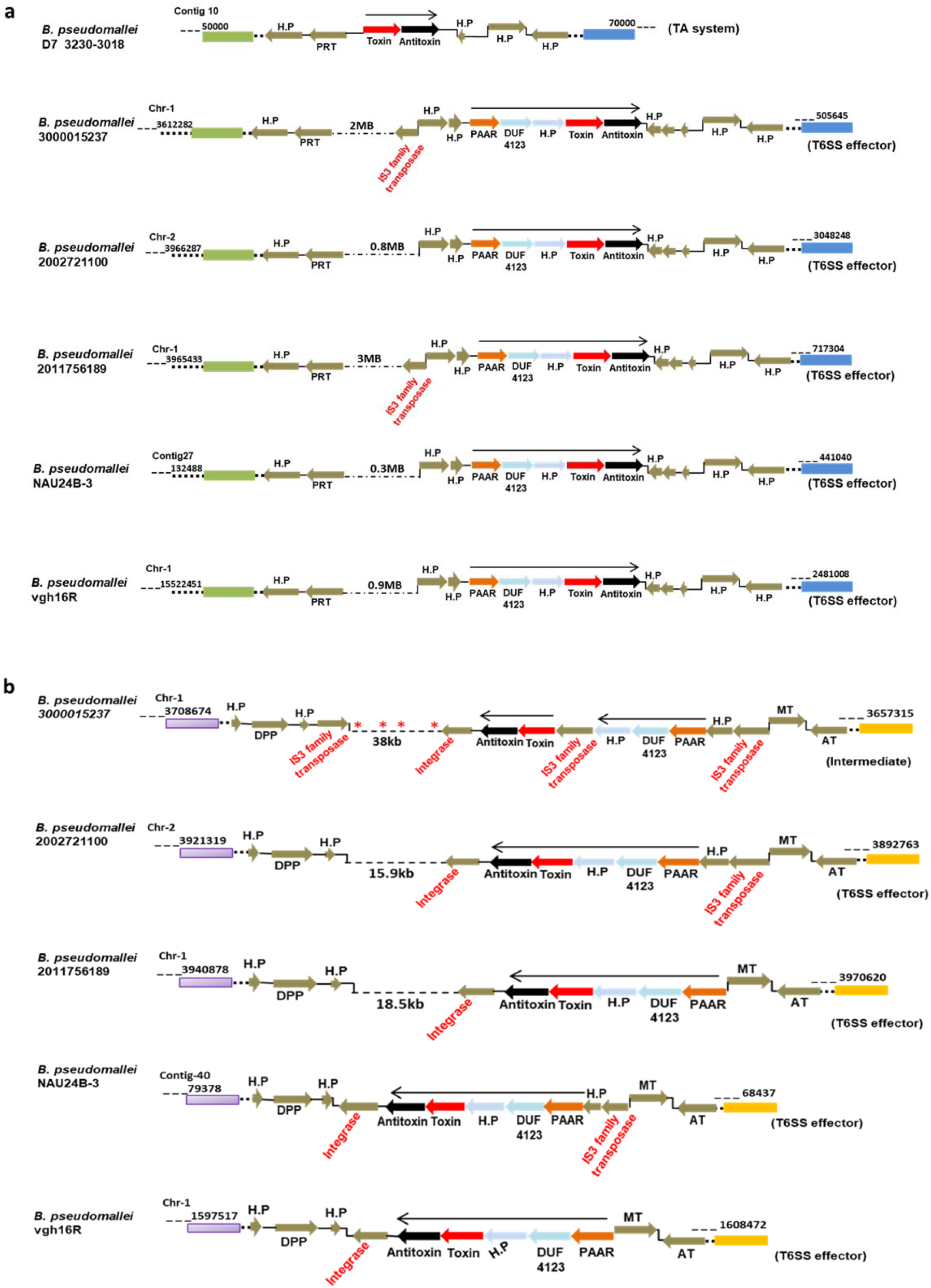
Conversion of Tse orthologs of a TA system into T6SS effectors in *B. pseudomallei*. (a) The genomic loci containing Tse orthologs (analogous to toxins) and cognate immunity protein (analogous to antitoxins) in different *B. pseudomallei* strains. The toxin and antitoxins were either encoded together (considered as TA) or encoded along with certain T6SS related (PAAR, DUF4123 and hypothetical protein) genes (considered as T6SS effector). The conservation of flanking genes reflects potential evolution occurring at the same genomic loci. (b) Intermediary stages of conversion of Tse orthologs into T6SS effectors in different *B. pseudomallei* strains. The sequential insertion and excision of IS3 transposable elements had potentially created operonic fusion of TA genes with the T6SS related genes, converting them into T6SS effector. Asterisks indicates presence of IS3 family transposase.

We further analysed whether the immunity protein (17tsi/38tsi) can regulate its own promoter. For this, the expression of *β-glucuronidase* reporter gene under the promoter of Tse effector-immunity operon was analysed in presence or absence of immunity protein (17tsi/38tsi) as well as effector-immunity pair (17tsei/38tsei). The reporter assays revealed that presence of Tsi/Tsei proteins significantly reduce the expression of *β-glucuronidase* (Fig. 4c and d).

### The Tse proteins are either encoded as T6SS effector or part of TA system in different bacteria

The Pfam database search revealed that homologs of the Tse and the Tsi proteins are conserved in diverse bacteria; predominantly being encoded together in an operon (Fig. 5 and Supplementary Fig. 7). As observed in the Tse effectors of NGJ1, presence of certain T6SS related proteins (PAAR, DUF4123 and hypothetical protein) was observed in the operons encoding Tse orthologs in certain Gamma-proteobacteria (class Morganellaceae and Pseudomonadaceae) and most of the Beta-proteobacteria (class Alcaligenaceae, Burkholderiaceae, Oxalobactereceae) (Supplementary Fig. 7). This suggested that these Tse orthologs may serve as T6SS effectors. However, some closely related strains of these bacteria as well as several Actinobacteria and Delta-proteobacteria were found to encode the Tse and Tsi orthologs in the operon and did not carry the T6SS associated PAAR, DUF4123 and hypothetical protein (Fig. 5 and Supplementary Fig. 7). The lack of T6SS related functions in the latter category of Tse and Tsi ortholog operons, suggested that they may not be secreted through T6SS and that they may be part of a toxin-antitoxin (TA) system. As indicated earlier, Tse proteins act as toxins and the Tsi proteins act as anti-toxins.

### The Tse proteins have potentially evolved as T6SS effectors from bacterial TA system

An examination of the genome of approximately 1800 strains of different *Burkholderia* sp. that are available in the Burkholderia genome database^36^, revealed intraspecific variation wherein some of them appear to have Tse orthologs as TA system while others appear to be T6SS effector (Supplementary Fig. 8). For example, the *B. pseudomallei* strain D7 3230-3018 harbours a Tse ortholog as a TA system while other *B. pseudomallei* strains encode the orthologs as a T6SS effector (at the same genomic locus) (Fig. 6a). The presence of many IS3 family transposases at the Tse locus suggested that they may have a role in generating this diversity. This was more apparent in *B. pseudomallei* strain 3000015237 wherein two IS3 elements carrying the T6SS related genes (PAAR, DUF4123 and Hypothetical protein) were present immediately upstream of TA genes (Fig. 6b). In another *B. pseudomallei* strain (NAU24B-3 and 2002721100); one of the two IS3 elements was absent. Moreover, in yet another *B. pseudomallei* strains (vgh16R and 2011756189), both the IS3 elements had been excised, leading to fusion of TA and T6SS related genes in a single operon (Fig. 6b). In *B. stagnalis* strains, the Tse orthologs are present as either a TA system or as a T6SS effector at the same genomic locus (Supplementary Fig. 9). This suggests that although we don’t yet have strains with intermediate genomic features, similar events involving IS elements may have possibly led to conversion of a TA system to a T6SS system in the *B. stagnalis*. Overall, our data supports that the Tse orthologs have evolved as T6SS effectors from ancestral TA system.

## Discussion

The genus Burkholderia constitutes a large group of bacterial species, being present as soil dwellers or living in association with plants, animals, fungi (endosymbiont) and insects (as symbiont)^37–39^. Recently, we had demonstrated that the rice associated *Burkholderia gladioli* strain NGJ1 utilizes a type III secretion system (T3SS) to feed on fungi (phenomenon known as mycophagy)^40^. In this study, we report that the NGJ1 bacterium not only has antifungal activity, but it also has anti-bacterial activity and that it utilizes two different type VI secretion systems (T6SS-1 and T6SS-2) to kill co-habiting bacteria.

The presence of a diverse arsenal of T6SS effectors highlights the potency of the antibacterial repertoire of NGJ1. To protect itself and sister cells from intoxication, the NGJ1 encodes an immunity protein that is specific for each of the effector proteins. This is evidence from the observation that co-expression of the immunity protein protects *E. coli* cells in which the cognate effector is ectopically expressed. A major exception was the observation that the Imm52 domain containing immunity (Tsi) proteins that are associated with Tox-REase5 domain (PF15648) containing effectors (Tse) failed to protect the cells (when co-expressed using two different plasmids). Notably when the effector and immunity proteins were expressed as transcriptionally fused proteins (Tsei), the cells survived. This suggests that the stoichiometric ratio of effector (Tse) and immunity (Tsi) proteins influences the effector neutralization ability of the immunity proteins. Additionally, the Tsi proteins possess a transcriptional repression activity, including ability to repress its own promoter. This suggests that the Tsi immunity protein neutralizes the cognate effector not only through direct interaction, but also through repression of transcription of the effector gene. It is interesting that the anti-toxin of certain toxin-antitoxin (TA) systems has been shown to have repressor activity^29,31^ and we note that the effector-immunity protein and the TA proteins are analogous to each other.

Recently, one of the Tox-REase-5 domain containing proteins (TseT) of *Pseudomonas aeruginosa* PAO1 has been shown to be a T6SS effector^23^. The *TseT* shares similar genomic organization to that of 17tse and 38tse of NGJ1, wherein certain T6SS related proteins (PAAR, DUF4123: a chaperone and Hypothetical protein: a co-chaperone) are encoded in the effector-immunity operon. These T6SS related proteins are shown to be required for T6SS mediated delivery of TseT in *P. aeruginosa* PAO1. It has been shown that the TseT effector interacts with the PAAR protein that is encoded as an upstream ORF in the same operon and that PAAR interacts with the VgrG of the T6SS apparatus^23^. In NGJ1, we observed a physical interaction of the Tse effector (17tse/ 38tse) protein with the PAAR protein that is encoded as upstream ORF in the same operon. Protein-protein interaction studies further suggested that the PAAR protein acts as a carrier of the Tse effectors and assists in their delivery by interacting with the VgrG of the T6SS-2 apparatus. This suggests that certain proteins such as PAAR, chaperone and co-chaperone that are encoded in the Tse-Tsi operon, are required for T6SS mediated delivery of Tse effector.

Bioinformatics analysis indicates that in Burkholderia and certain other bacteria, the Tse and Tsi orthologs are present in either one of two configurations. In one configuration they are encoded along with PAAR, chaperone and co-chaperone proteins, indicating that they may be part of T6SS effector-immunity pair. In the other configuration, only Tse and Tsi are present and the accessary proteins (PAAR, chaperone and co-chaperone) are absent. We think that in the second configuration, the Tse-Tsi proteins are part of a toxin-antitoxin (TA) system as they are analogous to toxins and antitoxin proteins in many bacteria.

This reflects an interesting scenario, wherein the Tse orthologs function as TA systems in certain bacteria and as T6SS effectors in other bacteria. The presence of Tse orthologs as TA in some strains of certain *Burkholderia* sp. and as T6SS effectors in other strains of the same species (at the same genomic locus), suggests that the Tse proteins might have evolved as T6SS effectors from an ancestral TA system. The IS3 family transposons seems to have played an important role in conversion of TA into T6SS effectors, by integrating T6SS related proteins (PAAR, DUF4123 and co-chaperon) at the upstream regions of putative TA operons (as observed in case of *B. pseudomallei* strain 3000015237). With sequential excision of IS3 elements, the T6SS related genes have become operonic with TA genes, leading to conversion of Tse orthologs of a TA system into T6SS effectors.

The toxins of the TA system predominantly function intracellularly and contribute towards bacterial growth arrest/ persistence/ biofilm formation^30,41,42^. By converting an intracellular toxin into an extracellular weapon that can be secreted through the T6SS, certain Burkholderia strains have gained the ability to inhibit growth of co-habiting bacteria. In NGJ1 it is apparent that the T6SS helps the bacterium in inhibiting the growth of a number of bacteria that co-habit the same niche. By inhibiting the growth of these co-habiting bacteria, NGJ1 may be reducing competition for resources including that may be released upon degradation of fungal biomass.

Overall, our study indicates that the NGJ1 bacterium uses two different T6SSs to inhibit growth of co-habiting bacteria. One category of potential T6SS effectors that contain Tox-REase-5 domain are related to the type II TA system. Analysis of the genomes of several Burkholderia strains suggests an ancestral TA system has been converted into a T6SS effector by a series of genetic recombination events involving IS3 elements. Thus as ancestral function involved in growth arrest/persistence appears to have been converted into an offensive weapon to inhibit growth of potential bacterial competitors.

## Materials and methods

### Growth conditions

The bacterium *Burkholderia gladioli* strain NGJ1 (NGJ2; rif^R^ derivative of NGJ1) and its derivative strains were grown on PDA (Potato Dextrose Agar; Himedia, India) plates at 28^°^C. *Escherichia coli* and its derivatives were grown on LBA (Luria Bertani Agar; Himedia, India) plates at 37^°^C. *Agrobacterium tumefaciens* strain EH101 was grown on PDA at 28^°^C. The *Saccharomyces cerevisiae* (yeast) strains were grown on YPD (Yeast Extract Peptone Dextrose; Himedia, India) at 28^°^C. Whenever required, the media was supplemented with antibiotics: Kanamycin, 50µg/ml; Rifampicin, 20µg/ml; Ampicillin, 50µg/ml and Spectinomycin, 50µg/ml. The list of NGJ1, *E. coli* and *S. cerevisiae* strains and various plasmids used in this study are summarized in Supplementary Table 4.

### Antibacterial activity

Various rice endophytic bacteria were isolated from field grown surface sterilized~45-day old Pusa Basmati-1 (PB1) rice leaves. List of various endophytic bacteria used in this study and their growth conditions are summarized in Supplementary Table 5. The pure cultures of each of these bacteria were established and 16s-ribosomal DNA sequencing was performed to identify them (primers are listed in Supplementary Table 6).

The antibacterial activity of NGJ1 and its different mutants was tested on solid laboratory media. 100 µl of overnight grown target bacteria were spread plated onto their respective growth medium (Supplementary Table 5) and 10 µl of overnight grown cultures of NGJ1 and its variants were spotted on the plate. The plates were incubated at ambient temperature (as per the target bacterium) and zone of inhibition was recorded 24h post incubation. The experiment was performed in triplicates and independently repeated three times. Similar results were observed in each biological and technical replicates.

### In-silico mining of T6SS apparatus and effector encoding gene clusters

The draft genome sequence of *B. gladioli* strain NGJ1^43^ was used for in-silico identification of putative T6SS apparatus encoding gene clusters using a web-based online tool; T346Hunter^44^. The presence of T6SS apparatus in the NGJ1 genome was further verified using Burkholderia Genome Database^36^. We carried out BLASTN analysis using T6SS apparatus components (such as HCP, VgrG and PAAR) and chaperone (DUF4123, DUF1795 and DUF2169) encoding genes of NGJ1 to predict T6SS effectors of NGJ1. The genomic organization of predicted T6SS effectors was analyzed using Burkholderia Genome Database and presence of cognate immunity proteins along with certain T6SS related proteins in each operon was noteworthy. The NCBI conserved domain database^45^ and Pfam database^46^ were used to identify conserved domain present in various T6SS effector and immunity proteins of NGJ1. For structure homology, SWISS-MODEL; an online protein structure similarity prediction tool^47^ was used.

### Construction of T6SS mutants

Partial fragments (~300 bp) of one of the core T6SS apparatus genes of T6SS-1 (*VipA*) and T6SS-2 (*ImpE*) were PCR amplified from the genomic DNA of *B. gladioli* strain NGJ1 using gene specific primers (Supplementary Table 6) and cloned into pK18 mob vector. The recombinant plasmid was electroporated (Gene pulsar XcellTm; BioRad) into *B. gladioli* strain NGJ1, as per the method described in^40^. The insertion mutants were selected on kanamycin and rifampicin containing KBA (King’s medium B Base; Himedia, India) plates. The ΔT6SS-1 and ΔT6SS-2 mutant NGJ1 strains were confirmed by PCR using gene specific flanking forward and vector specific reverse (M13) primers (Supplementary Table 6).

### Western blot analysis

The cell free supernatant was collected from 100ml of overnight grown NGJ1 culture and precipitated using TCA (12% wt/vol). The precipitated pellet was dissolved in 2 ml PBS (10mM) and used as crude supernatant protein. Further to isolate total protein, the bacterial pellet (obtained from 10 ml culture) was crushed in liquid N_2_ and the powder was dissolved in 2 ml of buffer (10 mM PBS: phosphate buffer saline; pH 7.4, 1 mM lysozyme, and 1 mM PMSF: Phenylmethanesulfonyl fluoride). Upon centrifugation, the soluble fraction was used as protein extract.

10 μg of protein samples were resolved on SDS-PAGE gel (12%) and electro blotted onto PVDF membrane, as described in^40^. The membrane was individually probed with HCP-1 (T6SS-1) and HCP-2 (T6SS-2) specific polyclonal peptide antibodies (primary antibody raised in rabbit) at 1:25,000 dilutions. The alkaline phosphatase conjugated anti-rabbit IgG (Sigma) was used as secondary antibody (1:10,000 dilutions) and the blot was developed as per manufacturer’s instruction (Sigma).

### Effector-immunity functionality assay

The full-length coding sequences of selected effector and immunity genes were PCR amplified from NGJ1 genomic DNA using gene specific primers (Supplementary Table 6). The effector encoding genes were cloned in pET23b (pET23b: effector) and the cognate immunity encoding genes were cloned in pET28a (pET28a: immunity). *E. coli* BL21 (DE3) cells were transformed with the pET23b: effector or co-transformed with the cognate pET28a: immunity plasmid. The positive transformed bacterial cells were selected by colony PCR using gene specific primer (Supplementary Table 6).

In order to test the toxicity of T6SS effectors, the recombinant *E. coli* cells were grown in 10 ml liquid media to mid log phase (O.D_600_ ~ 0.5) and the protein expression was induced using 1mM IPTG (Isopropyl β-d-1-thiogalactopyranoside; Sigma-Aldrich, USA) for 3h at 37°C. In control (-IPTG), an equivalent amount of sterile distilled H_2_O was added instead of IPTG. Subsequently, the recombinant *E. coli* cells were serially diluted and spotted on antibiotic containing LBA plates to monitor their growth. Similarly, the survival of *E. coli* cells that co-express effector and cognate immunity proteins were also analyzed.

Further the complete coding sequences of 17tse and 38tse effector along with their cognate immunity genes were PCR amplified from NGJ1 genomic DNA using effector specific forward and immunity specific reverse primers (Supplementary Table 6). The PCR product was cloned into pET28a to obtain pET28a:17tsei/pET28a:38tsei constructs and they were transformed into BL21 cells to express transcriptionally fused effector-immunity (17tsei/ 38tsei) proteins. Positive transformants were selected by colony PCR. Upon 3h of IPTG induction, the survival rate of the recombinant bacteria was analyzed as described above.

### Nuclease assay

For protein isolation, 10 ml cultures of recombinant *E. coli* cells (BL21) were grown to mid log phase (O.D_600_ ~ 0.5) and subsequently 1mM IPTG was added followed by incubation at 37°C for 3h. The bacterial pellet obtained upon centrifugation was sonicated in 1 ml buffer (10 mM PBS; pH 7.4, 1 mg/ml lysozyme and 1 mM PMSF). The soluble fraction was then collected and used as crude protein. Similarly, the crude protein was also isolated from non-recombinant BL21 cells as control to rule out the activity of other proteins present in the crude preparation.

1 μg of lambda DNA (Thermo Scientific™) and 0.5 μg of pET23b plasmids (linearized with *EcoR*I digestion) were treated with the 10 µl of different crude protein preparations for 30 min at 28°C. The buffer treated lambda DNA and linearized pET23b plasmid were used as control. The treated samples were resolved on 0.8% agarose gel and visualized under UV Tran-illuminator (ChemiDoc MP System, Bio-Rad).

The plasmid from various recombinant *E. coli* cells (10 ml culture) upon 3h of 1mM IPTG induction was extracted using geneJET plasmid isolation Kit (Thermo scientific). 100ng of isolated plasmids were transformed into chemical competent *E. coli* strain DH5α using heat-shock treatment. The positive transformants obtained on appropriate antibiotic containing LBA plates were counted.

Each experiment was independently repeated three times and similar results were obtained.

### Sytox Green staining and microscopic analysis

To visualize nucleic acid degradation under in-vivo condition, 10 ml of recombinant *E. coli* (BL21) cells were grown in LB media containing appropriate antibiotics. Upon 3h of 1 mM IPTG induction, 2 ml of culture was pelleted down and washed thrice with 50 mM Tris buffer (pH-7.0). Cells were heat-treated for 2 min at 90°C to make them permeable for uptake of the stain. The cells were stained with SYTOX™ Green Nucleic Acid Stain dye (0.2µM; Invitrogen, Catalog number: S7020) for 10 min in dark at 37°C. To visualize the nucleic acid, cells were washed twice with Tris buffer and analyzed by fluorescence microscope (AOBS TCS-SP5; LEICA, GERMANY) under GFP filter (495 nm). The experiment was independently repeated three times, each with a minimum of three technical repeats and similar results were obtained.

### Bacterial two-hybrid analysis

In order to understand the potential T6SS-1/T6SS-2 mediated secretion of Tse effectors, bacterial two-hybrid analysis was performed following established protocol^48^. Genes encoding the proteins of interest (X and Y) were PCR amplified from the NGJ1 genomic DNA using appropriate primers (Supplementary Table 6) and cloned into pKNT25 and pUT18C vectors, respectively. The recombinant plasmids encoding the T25-X and T18-Y hybrid proteins were transformed into the competent BTH101 reporter cells. The interaction was visually monitored by appearance of blue color due to activity of β-galactosidase on X-gal (20 mg/ml; 5-bromo-4-chloro-3-indolyl-β-D-galactopyranoside). Further, color formation was quantified spectrophotometrically by monitoring the absorbance at wavelength 450nm. The bacterial cells transformed with plasmids pKT25-zip and pUT18C-zip served as positive controls and those with pKNT25 and pUT18C served as negative controls.

### Yeast two hybrid analysis

Yeast two hybrid (Y2H) assay was performed using Matchmaker Gold Yeast Two-Hybrid library screening system (Clontech, USA). The full-length copy of genes of PAAR and Tse effector were PCR amplified from the NGJ1 genomic DNA using gene specific primer pairs (Supplementary Table 6) and cloned into pGBKT7 and pGADT7 vectors, respectively. Both the recombinant plasmids were co-transformed into Y2H Gold strain of yeast, following the manufacturer’s instructions. Positive transformants were selected on synthetically defined media lacking leucine and tryptophan amino acids (SD^−Leu-Trp^; double dropout). The interaction assay was performed by growing positive transformed colonies on synthetically defined media lacking leucine, tryptophan, histidine and adenine (SD^−Leu-Trp-His-Ade^; quadruple dropout) but supplemented with 30mM concentration of 3AT (3-Amino-1, 2, 4-triazole) at 30^0^ C. The yeast cells co-transformed with pGBKT7-53 and pGADT7-T were used as positive control while the cells co-transformed with pGBKT7 and pGADT7 (empty vectors) were used as negative control.

### Trans-repression assay

For trans-repression assay, the full-length coding sequence of genes of interest was PCR amplified from NGJ1 genomic DNA using gene specific primers (Supplementary Table 6). They were cloned in pGBKT7 vector (having both BD and AD domain) in frame with GAL4 transcription factor and transformed into *S. cerevisiae* (yeast) strain AH109 (harboring GAL4 regulated reporter genes lacZ; β-galactosidase, HIS3; imidazole glycerol phosphate dehydratase, and ADE2; Phosphoribosyl aminoimidazole carboxylase) as described^35^ using EZ-Yeast^TM^ Transformation Kit (MP Biomedicals). Positive colonies were selected through colony PCR using gene specific primers (Table 6) and the activity of reporter genes was tested by their differential growth on SD^**-**Trp-His-Ade^ (Triple dropout) media with 20 mM 3AT (3-amino-1, 2, 4-triazole). Further LacZ (β-galactosidase enzyme) expression was quantified spectrophotometrically (λ=450nm) using X-GAL as substrate.

### Auto trans-repression assay

The promoter region (500bp) of Tse effector operons (17tse and 38tse) was PCR amplified (Supplementary Table 6) and cloned into pBI101.1 expression vector in a manner that it can drive *GUS* (β-glucuronidase) expression. Recombinant pBI101.1 was transformed into *E. coli* BL21 (DE3) cells alone or along with pET28a: immunity (17tsi/38tsi/9tli) or pET28a: effector-immunity (17tsei/38tsei). Auto transcriptional repression activity was tested by visualizing the expression of *β-glucuronidase* gene using 5-bromo-4-chloro-3-indolyl glucuronide (X-Gluc) as substrate. Further, the β-glucuronidase enzyme activity was spectrophotometrically quantified by using a fluorescent substrate; 4-methylumbelliferyl ß-D-glucuronide (4-MUG)^49^.

### Phylogenetic analysis

The homologous amino acid sequences of Tse effectors and Tsi immunity proteins of NGJ1 were downloaded from Pfam^46^ and NCBI database^50^. After removal of duplicate sequences from the same strain, the Tse and Tsi homologous sequences were separately aligned using ClustalW algorithm. The aligned sequences were subjected to phylogenetic analysis using MEGAX following Neighbor-joining algorithm with 500 bootstrap values^51^.

### Evolutionary analysis

The genomic organization of Tse and Tsi orthologs were studied in different *Burkholderia* sp. using Burkholderia Genome Database^36^. In most of the strains these proteins were found to be encoded along with certain T6SS related proteins (PAAR, DUF4123: a chaperon and a hypothetical protein) and we considered them as candidate T6SS effectors. However, in some strains, only Tse and Tsi orthologs were encoded in the operon and the T6SS related proteins were absent. These orthologs were considered as part of Toxin-antitoxin (TA) system. Further the presence of IS3 family transposases were detected at these loci. The conservation of flanking genes was analyzed using NCBI blast analysis^50^ and Burkholderia Genome Database^36^.

### Statistical analysis

One-way analysis of variance was performed using Sigma Plot 12.0 (SPSS, Inc. Chicago, IL, USA) with P ≤ 0.001 and P ≤ 0.05 considered statistically significant. The statistical significance is mentioned in the figure legend, wherever required.

## Acknowledgements

SKY and JD acknowledge fellowship from DBT, Govt. of India. SG and RK acknowledge SPM and SRA fellowship from CSIR, Govt of India, respectively. We sincerely thank RV Sonti (NIPGR) for providing *E. coli* strains, Manjula Reddy (CSIR-CCMB) for sharing different strains/plasmids for bacterial two hybrid assay and Pinky Agarwal (NIPGR) for providing strains for yeast transcription repression assay. We also acknowledge Prabhu Patil and Kanika Bansal (IMTECH, Chandigarh) for valuable discussion and suggestions on genome analysis.

We sincerely thank RV Sonti, and SK Ray for valuable comments on the manuscript. This work was supported by core research grant from National Institute of Plant Genome Research, India. Also research funding from DBT, Government of India which support GJ lab is gratefully acknowledged. The funders had no role in study design, data collection and analysis, decision to publish, or preparation of the manuscript.

## Author contributions

GJ has overall planned the study and has supervised the experiments. SKY has initiated the project and handled most of the experiments presented in this study. AM and AK had assisted in characterization of T6SS effectors and their antibacterial property. SG carried out phylogenetic analysis and assisted in computational analysis. RK and JD had assisted in molecular cloning. SKY, AM, SG, and GJ contributed in manuscript writing and all authors had approved the manuscript.

