## Supplementary material for "Conversion of a defensive toxin-antitoxin system into an offensive T6SS effector in Burkholderia": Supplimentary Information

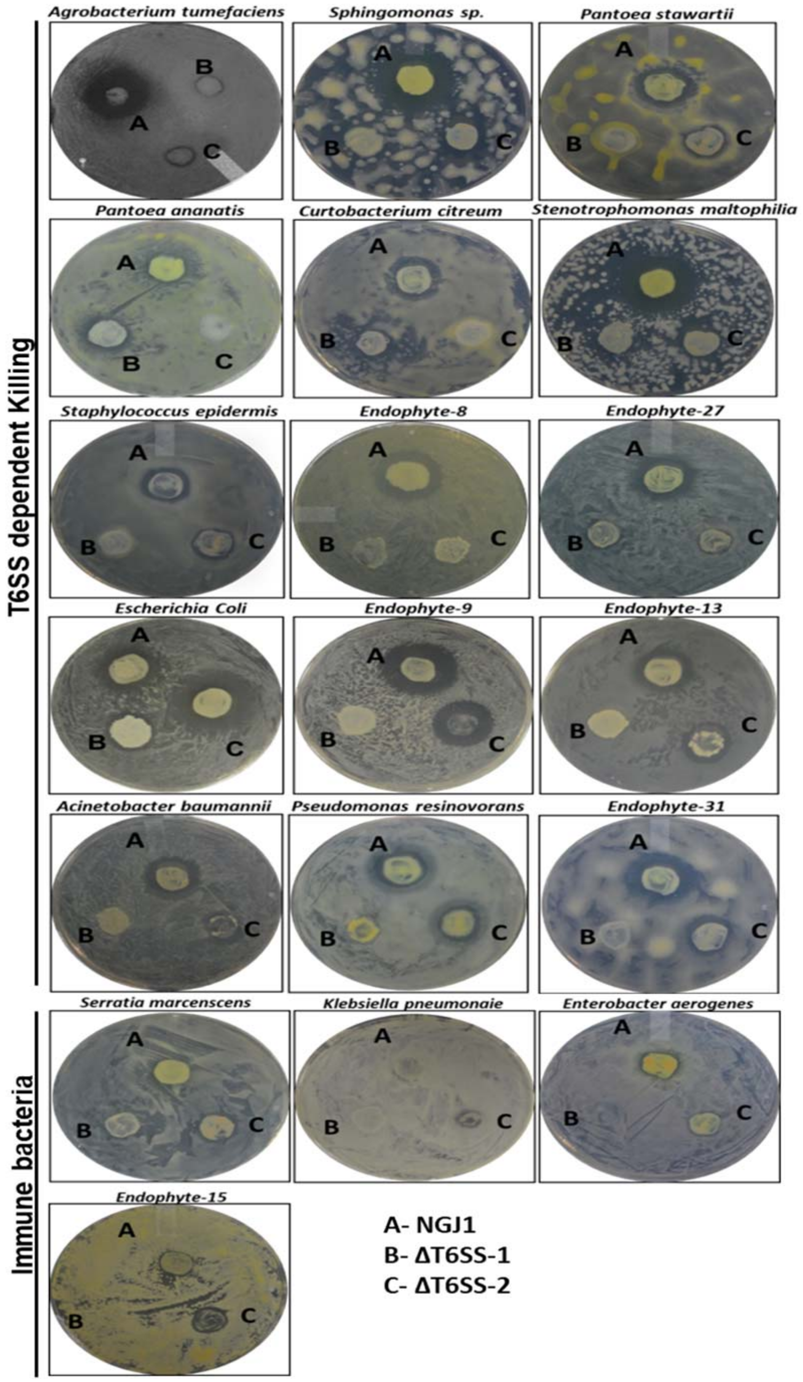

**Supplementary Fig. 1** ***B. gladioli*** **strain NGJ1 demonstrates broad spectrum antibacterial activity.** The antibacterial activity of NGJ1, ΔT6SS-1 and ΔT6SS-2 bacteria against various rice endophytes, *Escherichia coli* as well as *Agrobacterium tumefaciens* strains on solid media. The zone of inhibition reflected antibacterial activity of NGJ1 strains. Notably a few bacteria were found to be immune to NGJ1 strains.

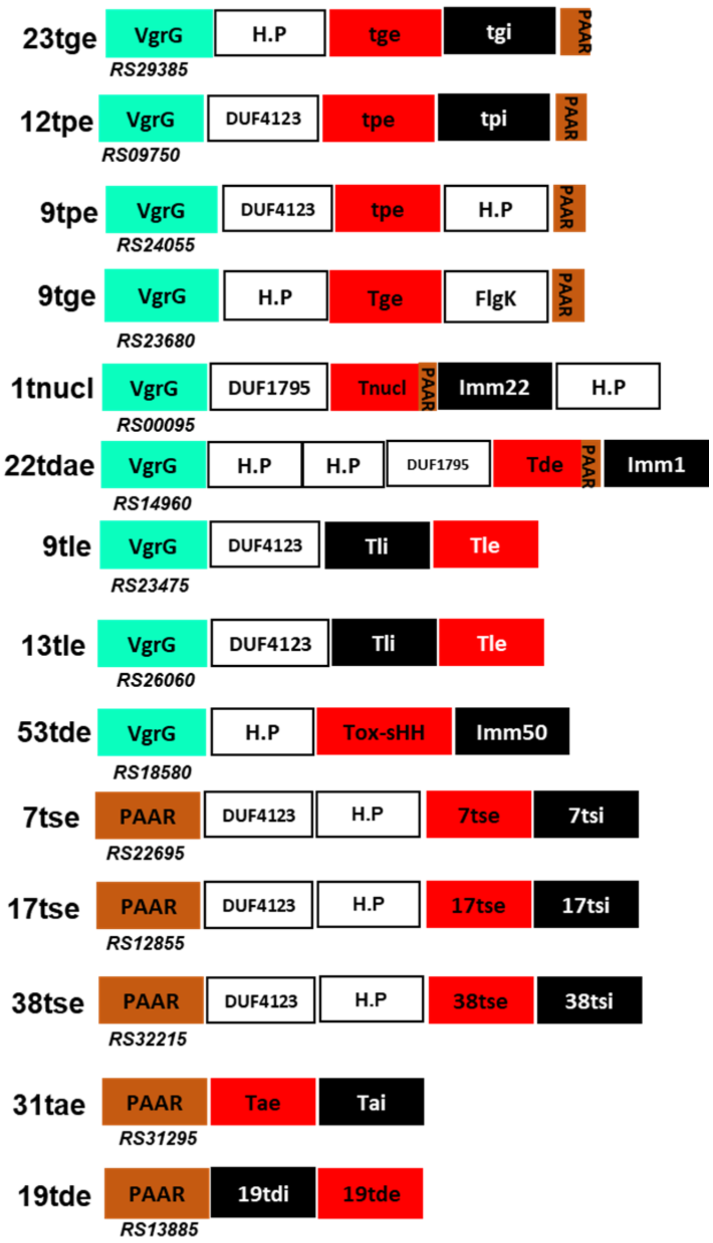

**Supplementary Fig. 2** ***B. gladioli*** **strain NGJ1 encodes various T6SS effectors.** Schematic representation of different T6SS effector operons present in the NGJ1 genome. PAAR genes were present as upstream ORF in some of the effector operon while in others the VgrG encoding gene was present at the upstream. Red and black color represents effector and immunity encoding gene, respectively. The name and the locus id as per Burkholderia genome database of each of the gene is mentioned.

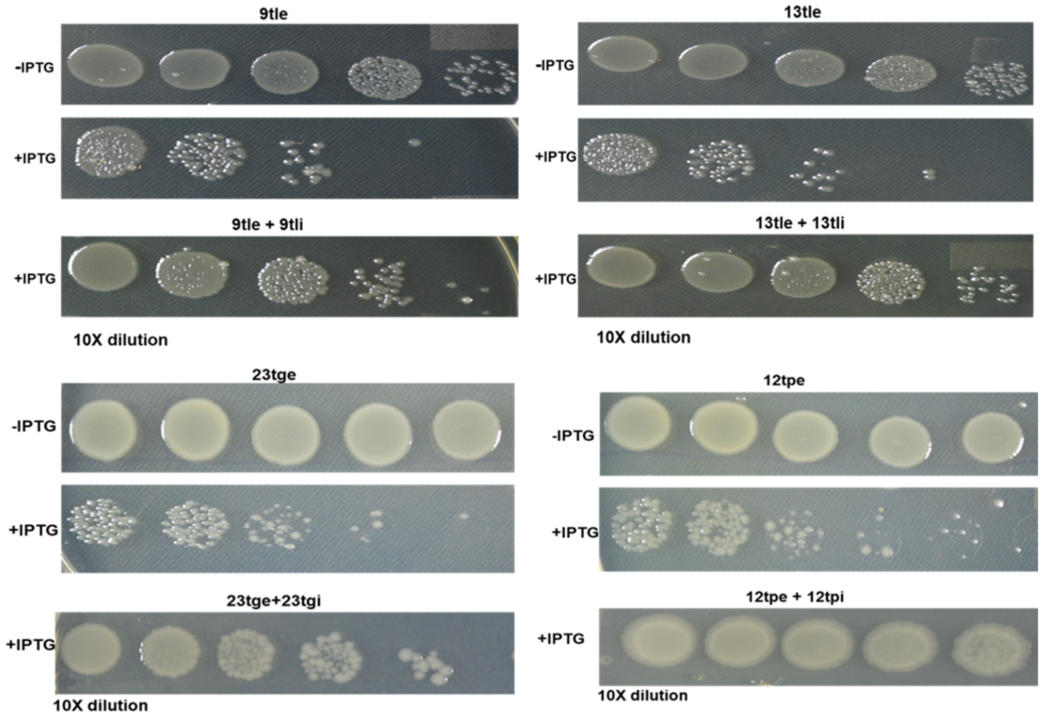

**Supplementary Fig. 3 The T6SS effectors of NGJ1 demonstrate antibacterial activity.** The ectopic expression of selected effectors (9tle, 13tle, 23tge, and 12tpe) inhibited the growth of *E. coli* (BL21) cells while co-expression with their cognate immunity protein (9tli, 13tli, 23tgi or 12tpe) using different plasmids protected the *E. coli* cells from effector mediated killing.

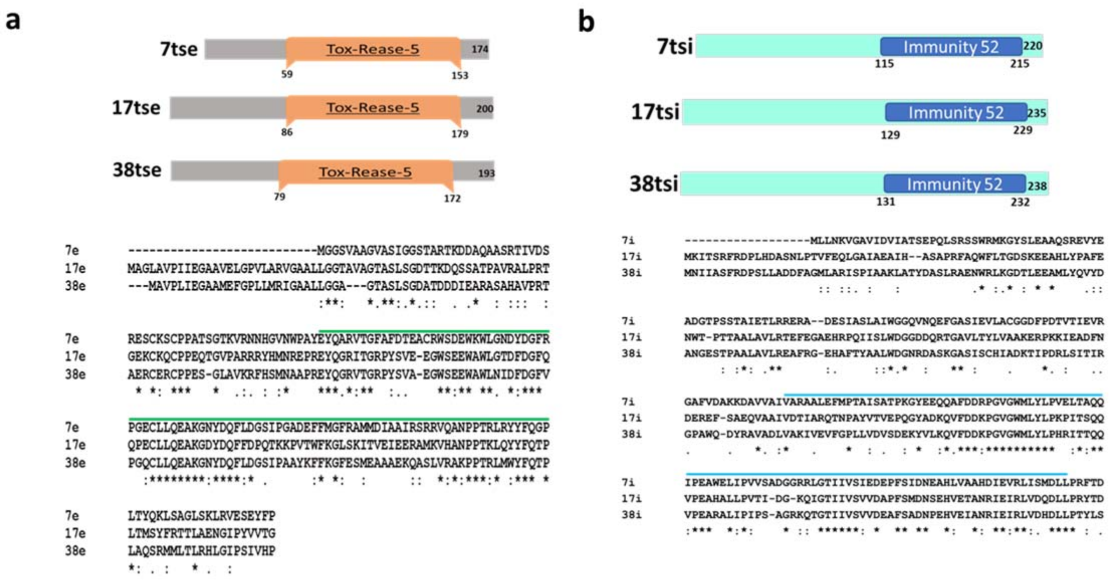

**Supplementary Fig. 4** Domain analysis showed presence of conserved (a) Restriction endonuclease fold toxin 5 (Tox-REase-5) in various Tse (7tse, 17tse and 38tse) effectors of NGJ1 and (b) Imm52 domain in the cognate Tsi immunity (7tsi, 17tsi and 38tsi) proteins. The multiple sequence alignment of these effector/ immunity proteins represents the sequence conservation. The presence of conserved domain is highlighted by color lines.

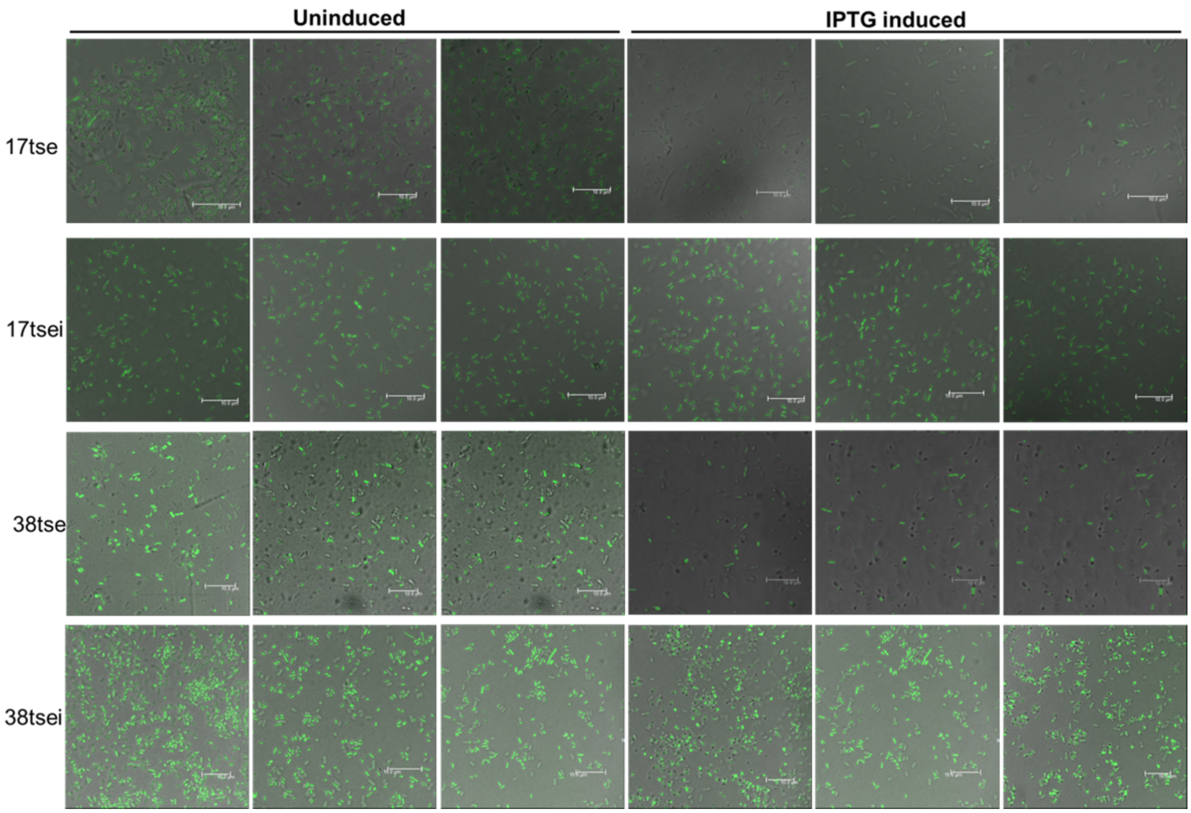

**Supplementary Fig. 5 Expression of Tse causes DNA degradation.** Fluorescence microscopic images of SYTOX-green (nucleic acid staining dye) stained *E. coli* cells that express either effector (17tse/38tse) or transcriptionally fused effector-immunity (17tsei/38tsei) proteins under IPTG induction (scale bar = 10μm). Lack of staining in the effector expressing cells suggested DNA degradation while proper staining in the effector-immunity (17tsei/38tsei) expressing cells suggested presence of intact DNA.

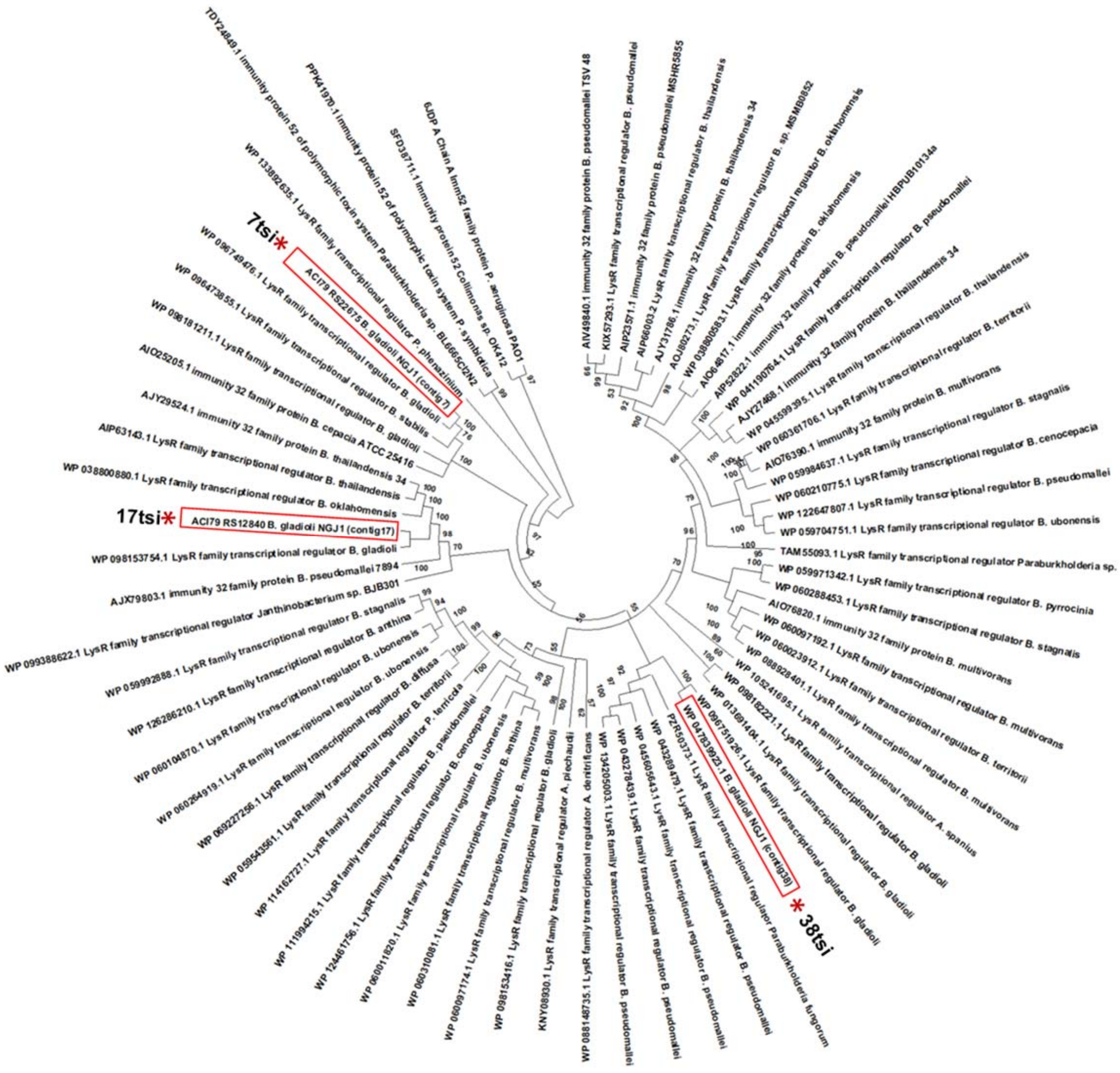

**Supplementary Fig. 6 The Tsi Immunity proteins of NGJ1 encodes LysR family transcription regulator.** The phylogenetic analysis was conducted using different orthologs (n=153) of Tsi immunity protein of different *Burkholderia* sp. The analysis reflected close phylogenetic relation of Tsi proteins of NGJ1 with bacterial LysR family of transcription regulator. The evolutionary history was inferred by MEGAX using Neighbor-Joining method. The values beside each node denote the bootstrap value. Asterisks and red boxes denote Tsi proteins of NGJ1.

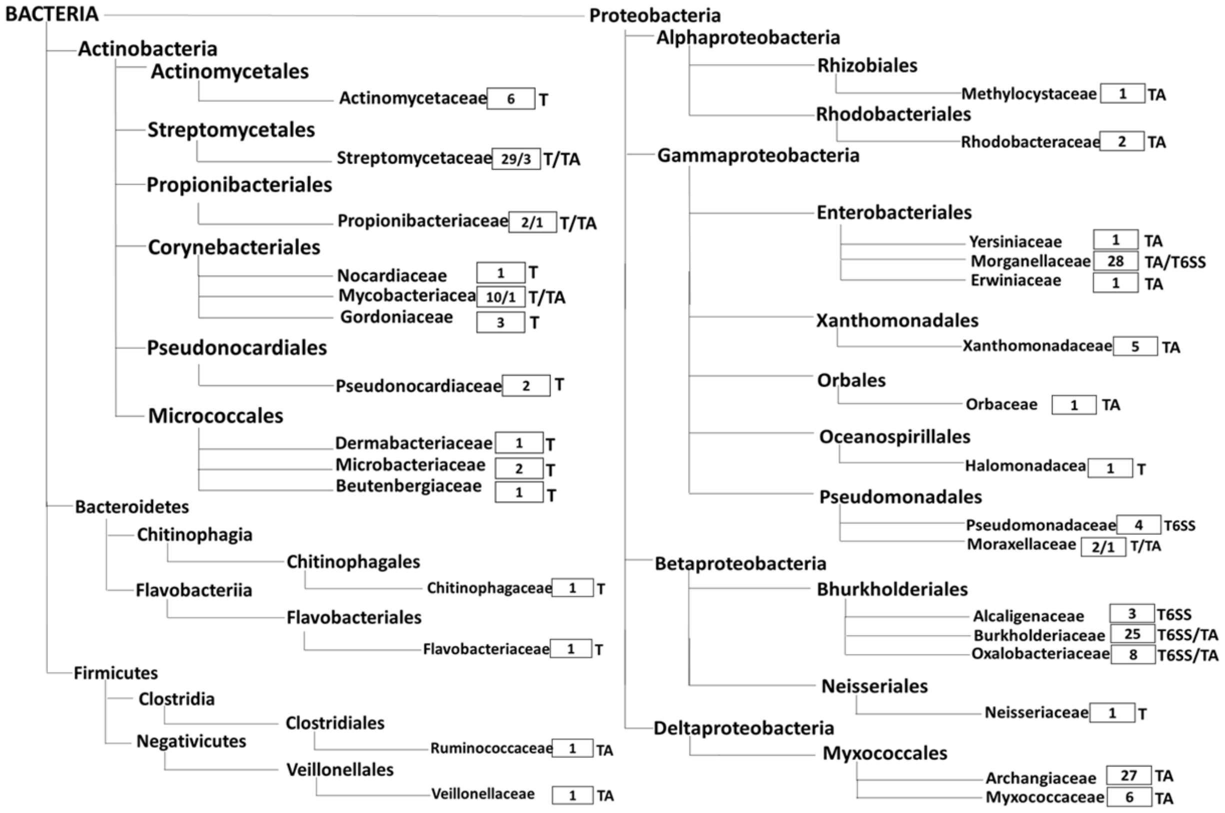

**Supplementary Fig. 7 The presence of Tse orthologs in different bacteria.** Pfam database reflected presence of Tse orthologs in different bacteria. The Tse orthologs were predominantly encoded as part of TA system or T6SS effectors. However, in a few bacteria, only the toxin (T) gene was found present, the immunity protein was absent in the operon. The number of T/TA/T6SS related Tse orthologs present in different bacterial family is reflected in empty box.

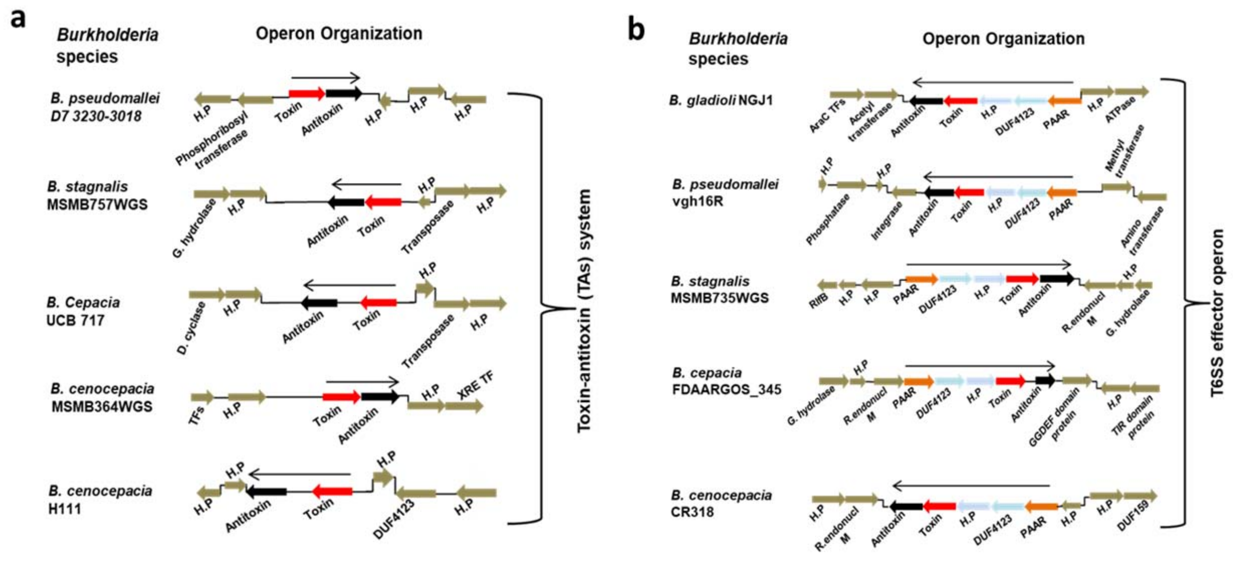

**Supplementary Fig. 8 Tse orthologs are encoded either as TA or T6SS effector in different *Burkholderia* sp.** The genomic loci encoding Tse orthologs as TA or T6SS effector in different *Burkholderia* sp are represented. Those having only toxin and antitoxin genes were considered as TA system while those having certain T6SS related genes [PAAR, DUF4123 and Hypothetical Protein (H.P)] in the operon, were considered as T6SS effector.

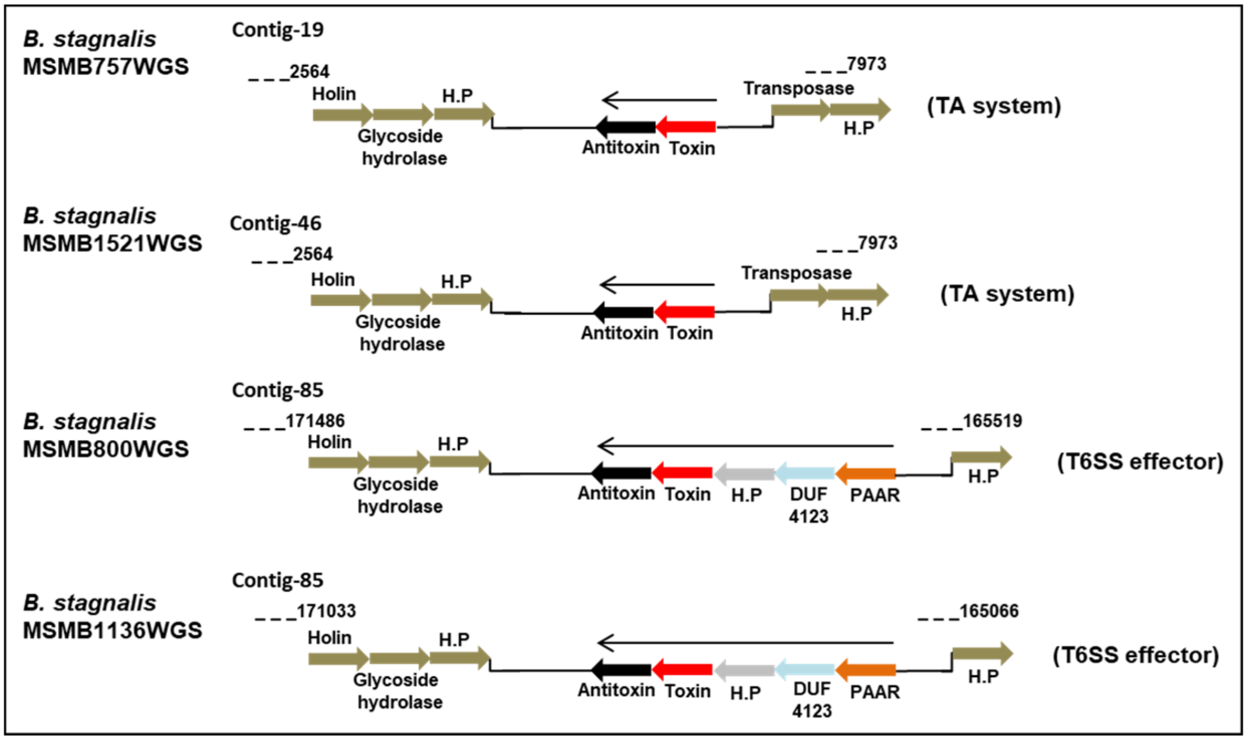

**Supplementary Fig. 9 The presence of Tse orthologs as TA or T6SS effector in *B. stagnalis***. The genomic loci encoding Tse orthologs as TA or T6SS effector in *B. stagnalis* strains are represented. The conservation of flanking genes reflected the conversion occurring at the same genomic loci.

**Supplementary Table 1 Antibacterial activity of NGJ1 against various endophytic bacteria**

| Endophytic bacteria | Antibacterial activity of NGJ1 | | |
|---|---|---|---|
| | Wild | ΔT6SS-1 | ΔT6SS-2 |
| *Escherichia coli* | ✓ | X | ✓ |
| *Agrobacterium tumefaciens* | ✓ | X | X |
| *Pantoea ananatis* | ✓ | X | X |
| *Pantoea stawartii* | ✓ | X | X |
| *Stenotrophomonas maltophilia* | ✓ | X | X |
| *Sphingomonas sp.* | ✓ | X | X |
| *Staphylococcus epidermis* | ✓ | X | X |
| *Pseudomonas resinovorans* | ✓ | X | ✓ |
| *Curtobacterium citreum* | ✓ | X | X |
| *Acinetobacter baumannii* | ✓ | X | X |
| *Endophyte-8* | ✓ | X | X |
| *Endophyte-9* | ✓ | X | ✓ |
| *Endophyte-13* | ✓ | X | ✓ |
| *Endophyte-27* | ✓ | X | X |
| *Endophyte-31* | ✓ | X | X |
| *Serratia marcenscens* | X | X | X |
| *Klebsiella pneumonaie* | X | X | X |
| *Enterobacter aerogenes* | X | X | X |
| *Endophyte-15* | X | X | X |

**Supplementary Table 2 T6SS effector repertoire of NGJ1 and their predicted functions**

| Effector | Motif | Effector ID | Target | |
|---|---|---|---|---|
| 9tle1 | Lipase | ACI79_RS23485 | Cell membrane | 75-90 |
| 13tle1 | Lipase | ACI79_RS26075 | Cell membrane | 60-75 |
| 31tae4 | Amidase | ACI79_RS31295 | Cell Wall | 300-285 |
| 7tse | Tox-REase-5 | ACI79_RS22680 | DNA | 95-75 |
| 17tse | Tox-REase-5 | ACI79_RS12845 | DNA | 55-35 |
| 38tse | Tox-REase-5 | ACI79_RS32205 | DNA | 215-200 |
| 1tnucl | tRNA Nuclease | ACI79_RS00105 | Nucleic acid | 95-115 |
| 22tdae | SCP_Deaminase | ACI79_RS14940 | Nucleic acid | 960-930 |
| 19tde | Nuclease | ACI79_RS13880 | Nucleic acid | 96-75 |
| 53tde | toxin_SHH | ACI79_RS18585 | Nucleic acid | 80-95 |
| 9tge | Muramidase | ACI79_RS23690 | Cell Wall | 680-700 |
| 9tpe | Aminopeptidase | ACI79_RS24045 | Proteins | 55-35 |
| 23tge | Glucosaminidase | ACI79_RS29395 | Cell Wall | 385-410 |
| 12tpe | Metallopeptidase | ACI79_RS09760 | Proteins | 50-70 |

**Supplementary Table 3 The structural similarity of Tsi immunity protein with CodY protein, using SWISS-MODEL**

| Template | Sequence identity | QSQE | Found by | Sequence similarity | Coverage | Description |
|---|---|---|---|---|---|---|
| rho.1.A | 50.64 | - | BLAST | 0.43 | 1 | Uncharacterized protein |
| 4rho.1.A | 50.43 | - | HHblits | 0.42 | 1 | Uncharacterized protein |
| 6jdp.1.A | 23.86 | - | BLAST | 0.34 | 0.84 | Imm52 family protein |
| 5ey2.1.B | 20.59 | 0.04 | HHblits | 0.33 | 0.14 | Pleiotropic repressor CodY |
| 5ey2.1.A | 20.59 | 0.04 | HHblits | 0.33 | 0.14 | Pleiotropic repressor CodY |
| 2gx5.1.A | 25.71 | 0.1 | HHblits | 0.34 | 0.15 | Pleiotropic repressor CodY |
| 5n0l.1.A | 22.86 | - | HHblits | 0.34 | 0.15 | Pleiotropic repressor CodY |
| 5n0l.1.F | 22.86 | - | HHblits | 0.34 | 0.15 | Pleiotropic repressor CodY |
| 5ey1.1.A | 22.86 | 0.06 | HHblits | 0.33 | 0.15 | Pleiotropic repressor CodY |
| 5ey0.1.A | 22.86 | 0.06 | HHblits | 0.33 | 0.15 | Pleiotropic repressor CodY |
| 5ey0.1.B | 22.86 | 0.06 | HHblits | 0.33 | 0.15 | Pleiotropic repressor CodY |
| 2hgv.1.A | 25.71 | - | HHblits | 0.34 | 0.15 | Pleiotropic repressor CodY |
| 5lnh.3.B | 26.47 | 0.22 | HHblits | 0.35 | 0.14 | Pleiotropic repressor CodY |
| 5lnh.1.B | 26.47 | 0.22 | HHblits | 0.35 | 0.14 | Pleiotropic repressor CodY |
| 5lnh.1.A | 26.47 | 0.22 | HHblits | 0.35 | 0.14 | Pleiotropic repressor CodY |
| 5lnh.4.A | 26.47 | 0.21 | HHblits | 0.35 | 0.14 | Pleiotropic repressor CodY |
| 5loo.1.A | 26.47 | - | HHblits | 0.35 | 0.14 | Pleiotropic repressor CodY |
| 5loo.1.B | 26.47 | - | HHblits | 0.35 | 0.14 | Pleiotropic repressor CodY |
| 5ey2.1.D | 20.59 | 0.03 | HHblits | 0.33 | 0.14 | Pleiotropic repressor CodY |
| 5ey2.1.C | 20.59 | 0.03 | HHblits | 0.33 | 0.14 | Pleiotropic repressor CodY |

### Supplementary Table 4 Bacterial strains and plasmids used in this study

| Strains or plasmid | Relevant characteristics | Source/reference |
|---|---|---|
| ***E. coli* strains** | | |
| DH5α | F', endA1 hsdR17 (rk- mk+) supE44 thi-1 recA1 gyrA TelA1 cp8OdlacZAM15 A (lacZY A-argF) U169 | Lab collection |
| BL21 | F- ompT gal dcm lon hsdSB(rB- mB-) λ(DE3 [lacI lacUV5-T7 gene 1 ind1 sam7 nin5]) | ,, |
| BTH101 | Non-reverting adenylate cyclase deficient (cya) *E. coli* reporter strain for bacterial two-hybrid system | Gift from Dr Manjula Reddy, CCMB, Hyderabad |
| ***B. gladioli* strains** | | |
| NGJ1 | Natural isolate | Lab collection |
| NGJ-2 | *rif*-2, $Rif^R$ derivative of NGJ-1 | ,, |
| ΔT6SS-1 | *VipA*::pK18mob, $Km^r$ derivative of NGJ-2 (T6SS-1 mutant strain) | Current study |
| ΔT6SS-2 | *ImpE*::pK18mob, $Km^r$ derivative of NGJ-2 (T6SS-2 mutant strain) | ,, |
| **Yeast strains** | | |
| Y2HGold | Strain for Yeast Two-Hybrid System; MATa, trp1-901, leu2-3, 112, ura3-52, his3-200, gal4Δ, gal80Δ, LYS2:: GAL1UAS–Gal1TATA–His3, GAL2UAS–Gal2TATA–Ade2, URA3:: MEL1UAS–Mel1TATA, AUR1-C, MEL1. | Lab collection |
| AH109 | Strain for Trans-repression assay; MATa, trp1–901, leu2–3, 112, ura3–52, his3–200, GAL4D, GAL80D, LYS2:: GAL1-UAS-GAL1TATA-his3, GALUASGAL2TATA-ade2 and URA3:: MEL1UAS-MEL1TATA-lacZ. | ,, |
| **Plasmids** | | |
| pK18mob | pUC18 derivative; Mob+ Tra– KanR | Lab collection |
| pET28a | $Kan^R$, Expression vector | ,, |
| pET23b | $Amp^R$, Expression vector | ,, |
| pK18mob::*VipA* | $Kan^R$, pK18mob harboring partial fragment of *VipA* | Current study |
| pK18mob::*ImpE* | KanR, pK18mob harboring partial fragment of *ImpE* | ,, |
| pET23b::17tse | $Amp^R$,pET23b harboring full length *17tse* | ,, |
| pET23b::*38tse* | $Amp^R$,pET23b harboring full length *38tse* | ,, |
| pET23b::*9tle* | $Amp^R$,pET23b harboring full length *9tle* | ,, |
| pET23b::*13tle* | $Amp^R$,pET23b harboring full length *13tle* | ,, |
| pET23b::*23tge* | $Amp^R$,pET23b harboring full length *23tae* | ,, |
| pET23b::*12mpe* | $Amp^R$,pET23b harboring full length *12mpe* | ,, |
| pET28a::*17tsi* | $Kan^R$, pET28a harboring full length *17tsi* | ,, |
| pET28a::*38tsi* | $Kan^R$, pET28a harboring full length *38tsi* | ,, |
| pET28a::*9tli* | $Kan^R$, pET28a harboring full length *9tli* | ,, |

| | | |
|---|---|---|
| pET28a::*13tli* | $Kan^R$, pET28a harboring full length *13tli* | ,, |
| pET28a::*23tgi* | $Kan^R$, pET28a harboring full length *23tai* | ,, |
| pET28a::*12mpi* | $Kan^R$, pET28a harboring full length *12mpi* | ,, |
| pET28a::*17tsei* | $Kan^R$, pET28a harboring full length *17tsei* (Fused ) | ,, |
| pET28a::*38tsei* | $Kan^R$, pET28a harboring full length *38tsei* (Fused ) | ,, |
| pKNT25 | $Kan^R$, Vector for Bacterial two-hybrid system | Gift from Dr Manjula Reddy, CCMB, Hyderabad |
| pUT18c | $Amp^R$, Vector for Bacterial two-hybrid system | ,, |
| pKNT25-zip | $Kan^R$, Bacterial two-hybrid system control plasmid | ,, |
| pUT18c-zip | $Amp^R$, Bacterial two-hybrid system control plasmid | ,, |
| pKNT25::*VgrG* (T6SS-2) | $Kan^R$, pKNT25 harboring *VgrG* of T6SS-2 operon | Current study |
| pKNT25::*VgrG* (9tle) | $Kan^R$, pKNT25 harboring *VgrG* of 9tle operon | ,, |
| pKNT25::*VgrG* (12tpe) | $Kan^R$, pKNT25 harboring *VgrG* of 12tpe operon | ,, |
| pUT18c::*Hcp* (T6SS-1) | $Amp^R$, pUT18c harboring *Hcp* of T6SS-1 operon | ,, |
| pUT18c::*Hcp* (T6SS-2) | $Amp^R$, pUT18c harboring *Hcp* of T6SS-2 operon | ,, |
| pUT18c::*PAAR* (17tse) | $Amp^R$, pUT18c harboring *Hcp* of 17tse operon | ,, |
| pUT18c::*PAAR* (38tse) | $Amp^R$, pUT18c harboring *Hcp* of 38tse operon | ,, |
| pGBKT7 | $Kan^R$, Yeast two-hybrid bait expression vector that is designed to express a fusion protein of the GAL4 DNA-binding domain (DNA-BD) | Lab collection |
| pGADT7 | $Amp^R$, Yeast two-hybrid prey expression vector that is designed to express a fusion protein of the GAL4 activation domain (DNA-AD) | ,, |
| pGBKT7::*PAAR* (17tse) | $Kan^R$, pGBKT7 harboring *PAAR* of 17tse operon | Current study |
| pGBKT7::*PAAR* (38tse) | $Kan^R$, pGBKT7 harboring *PAAR* of 38tse operon | ,, |
| pGBKT7::*17tse* | $Amp^R$, pGBKT7 harboring *17tse* | ,, |
| pGBKT7::*38tse* | $Amp^R$, pGBKT7 harboring *38tse* | ,, |
| pBI101.1 | $Kan^R$, Promoter less beta-glucuronidase gene (*GUS*) expressing vector | Lab collection |
| pBI101.1::*p17* | pBI101.1 harboring promoter of 17tse operon (500bp) | Current study |
| pBI101.1::*p38* | pBI101.1 harboring promoter of 38tse operon (500bp) | ,, |
| AD66 | $Kan^R$, pGBKT7 derived vector in which Activation domain (AD) of *GAL4* TF is cloned. | Gift from Dr Pinky Agarwal, NIPGR, N. Delhi |
| AD66::*17tsi* | $Kan^R$, AD66 harboring *17tsi* in fusion with *GAL4* TF | Current study |
| AD66::*38tsi* | $Kan^R$, AD66 harboring *38tsi* in fusion with *GAL4* TF | ,, |
| AD66::*9tli* | $Kan^R$, AD66 harboring *9tli* in fusion with *GAL4* TF | ,, |

**Supplementary Table 5 Growth conditions of various endophytic bacteria used during the study**

| **Bacteria** | **Growth Temperature** | **Growth media** |
|---|---|---|
| *Pantoea ananatis* | $28^0$ C | PDB/PDA |
| *Pantoea stawartii* | " | " |
| *Stenotrophomonas maltophilia* | " | " |
| *Sphingomonas sp.* | " | " |
| *Staphylococcus epidermis* | " | " |
| *Pseudomonas resinovorans* | " | " |
| *Curtobacterium citreum* | " | " |
| *Acinetobacter baumannii* | $37^0$ C | LB/LBA |
| *Endophyte-8* | " | " |
| *Endophyte-9* | " | " |
| *Endophyte-13* | " | " |
| *Endophyte-27* | $28^0$ C | PDB/PDA |
| *Endophyte-31* | " | " |
| *Serratia marcenscens* | " | " |
| *Klebsiella pneumonaie* | $37^0$ C | LB/LBA |
| *Enterobacter aerogenes* | $28^0$ C | PDB/PDA |
| *Endophyte-15* | $37^0$ C | LB/LBA |

### Supplementary Table 6 List of primers used in this study

| Primer name | Primer sequence |
|---|---|
| **T6SS mutant study** | |
| ΔVipA | Forward- ATAAGCTTATGGCGAAAAAAGAAAGTATCCAG |
| | Reverse- ATTGAATTCTTATTCGCCCTCTTTGTCGCC |
| ΔImpE | Forward- ATAAGCTTATGGATCCCCGCCTGCTTCGT |
| | Reverse- ATTGAATTCCTACAGGATCGGGCGCTT |
| M13 | Forward- TGTAAAACGACGGCCAGT |
| | Reverse- AGGAAACAGCTATGACCAT |
| 16s rDNA | Forward- GAGTTTGATCMTGGCTCAG |
| | Reverse- AAGGAGGTGATCCAGCC |
| **Functionality assay in *E.coli*** | |
| 17tse | Forward- CATATGGCGGGACTGGCAGTG |
| | Reverse- AAGCTTTCAACCAGTGACGACGTA |
| 38tse | Forward- CATATGGCAGTACCTCTCATCGAAG |
| | Reverse- AAGCTTTCAAGGGTGGACGATTGACGGT |
| 9tle | Forward- AAGCTTATGTCTATTAGACAAGCTGGGGCGC |
| | Reverse- CTCGAGTCAAGCGACAGCCGCGTCGT |
| 13tle | Forward- CATATGAGCAACGCGCCCATCATTC |
| | Reverse- AAGCTTTCATGCCGCCGATGCCGAGT |
| 23tge | Forward- GAATTCGTGGGACTGGCCTGTCGCGCGT |
| | Reverse- GCGGCCGCCTAACGGCTCCTGTCGATTTC |
| 12mpe | Forward- GTCGACATGGCAACCATCCCGGGCTGT |
| | Reverse- CTCGAGTCAAGCCCCCATTGCGGAATTGAA |
| 17tsi | Forward- GAATTCATGAAAATCACTTCCCGC |
| | Reverse- AAGCTTTCAGATGTCCGTGTATCTCG |
| 38tsi | Forward- GAATTCATGAACATCATCGCAAGCTTCG |
| | Reverse- CTCGAGTTATAAGCTGAGATAGGTCGGC |
| 9tli | Forward- GAATTCATGCGGTATTGGTTGGTGGG |
| | Reverse- AAGCTTTTACTTGCTCATGCAACCCG |
| 13tli | Forward- GAATTCATGACACCGAATACGAAACACG |
| | Reverse- CTCGAGTCAATTCCACTGAACCGGT |
| 23tgi | Forward- GGATCCATGAGCCAACACCATTCAT |
| | Reverse- GCGGCCGCCTATTCCTCTAGATCGTATTGAGTG |
| 12mpi | Forward- CATATGGACGAGCGCTTGATGAAG |
| | Reverse- AAGCTTTCATGTTGAGTCTCGTTGCG |
| **Bacterial two hybrid analysis** | |
| VgrG (T6SS-2) | Forward- AATTTCGGATCCCATGGCTCACGCAATCACGCTC |
| | Reverse- AATTTCGAATTCTCCAGGATCATCGGCGC |
| VgrG (9tle) | Forward- AATTTCGGATCCCATGACTCCCGACCACACCCT |
| | Reverse- AATTTCGAATTCTCGTTCAGATCGATCATGGTGGC |

| | |
|---|---|
| VgrG (12tpe) | Forward- AATTTCGGATCCCATGCAGAACACGGCACAAG |
| | Reverse- AATTTCGAATTCTCGGAGAGCGCGACAGAC |
| HCP (T6SS-1) | Forward- AATTTCGGATCCCATGTTAGATATCTATCTGAATTTCGGG |
| | Reverse- AATTTCGAATTCGACCGCGTAGGTCTTGTCGTT |
| HCP (T6SS-2) | Forward- AATTTCGGATCCCATGGCATTCGACATGCACCTG |
| | Reverse- AATTTCGAATTCCTGCTTGGCGACCTG |
| PAAR (17tse) | Forward- AATTTCGGATCCCATGAGCCGAGCCATGATCTGC |
| | Reverse- AATTTCGAATTCTCCGCGCGCAATCATTGT |
| PAAR (38tse) | Forward- AATTTCGGATCCCATGGCTATACGAGCGATCATTTGC |
| | Reverse- AATTTCGAATTCACCCCGGACGATCAT |
| **Yeast two hybrid analysis** | |
| 17tse | Forward- GAATTCATGGCGGGACTGGCAGTG |
| | Reverse- GGATCCTCAACCAGTGACGACGTA |
| 38tse | Forward- GAATTCATGGCAGTACCTCTCATCGAAG |
| | Reverse- GGATCCTCAAGGGTGGACGATTGACGGT |
| PAAR (17tse) | Forward- GAATTCATGAGCCGAGCCATGATCTGC |
| | Reverse- GGATCCCTATCCGCGCGCAATCATTGT |
| PAAR (38tse) | Forward- GAATTCATGGCTATACGAGCGATCATTTGC |
| | Reverse- GGATCCTCAACCCCGGACGATCAT |
| **Trans-repression assay** | |
| 17tsi | Forward- GAATTCATGAAAATCACTTCCCGC |
| | Reverse- GTCGACTCAGATGTCCGTGTATCTCG |
| 38tsi | Forward- GAATTCATGAACATCATCGCAAGCTTCG |
| | Reverse- CTGCAGTTATAAGCTGAGATAGGTCGGC |
| 9tli | Forward- GAATTCATGCGGTATTGGTTGGTGGG |
| | Reverse- GTCGACTTACTTGCTCATGCAACCCG |
| P17 (promoter) | Forward- AATTCAGACTCTCTTCCCCGACAATGCCAATGCGA |
| | Reverse- GTTACCTCTAGAGTGCTCTCTCGTAGGTTTGATTTCGT |
| P38 (promoter) | Forward- AATTCAGACTCAGAATCGCCGAAGGATGCCG |
| | Reverse- GTTACCTCTAGATTGATCCTCCCTTTGCTTGGTGCTC |
